# ELMER v.2: An R/Bioconductor package to reconstruct gene regulatory networks from DNA methylation and transcriptome profiles

**DOI:** 10.1101/148726

**Authors:** Tiago C Silva, Simon G Coetzee, Lijing Yao, Nicole Gull, Dennis J Hazelett, Houtan Noushmehr, De-Chen Lin, Benjamin P Berman

## Abstract

**Motivation:** DNA methylation has been used to identify functional changes at transcriptional enhancers and other cis-regulatory modules (CRMs) in tumors and other disease tissues. Our R/Bioconductor package *ELMER* (Enhancer Linking by Methylation/Expression Relationships) provides a systematic approach that reconstructs altered gene regulatory networks (GRNs) by combining enhancer methylation and gene expression data derived from the same sample set.

**Results:** We present a completely revised version 2 of *ELMER* that provides numerous new features including an optional web-based interface and a new Supervised Analysis mode to use pre-defined sample groupings. We show that this approach can identify GRNs associated with many new Master Regulators including *KLF5* in breast cancer.

**Availability:** *ELMER* v.2 is available as an R/Bioconductor package at http://bioconductor.org/packages/ELMER/

## 1 Introduction

Motivated by the identification of transcription factor binding sites (TFBSs), enhancers, and other cis-regulatory modules (CRMs) from DNA methylation data in tumor samples (Berman et al., 2012; Hovestadt et al., 2014; Johann et al., 2016), and the strong association between DNA methylation and target gene expression in tumors (Aran et al., 2013; Aran and Hellman, 2013), we previously developed an R/Bioconductor package *ELMER* (Enhancer Linking by Methylation/Expression Relationships) to infer regulatory element landscapes and GRNs from cancer methylomes (Yao et al., 2015). ELMER version 1 has been adopted by other groups (Dhingra et al., 2017; Mishra and Guda, 2017; Malta et al., 2018), and remains the only publicly available software tool to use matched DNA methylation and expression profiles to reconstruct TF networks (reviewed in Teschendorff and Relton, 2018). Other tools such as TENET (Rhie, 2016) and RegNetDriver (Dhingra et al., 2017) have incorporated ELMER principles and code into cancer network analysis.

We present here a substantially re-written ELMER v. 2 (Fig. 1A) that implements new features and improvements including: (i) support for Infinium HM450 or EPIC arrays and RNA-seq using the gold-standard MultiAssayExperiment (MAE) data structure, (ii) integration with our TCGABiolinks package (Colaprico et al., 2015) for cohort selection and data importing from the NCI Genomic Data Commons (Grossman et al., 2016), (iii) integration with our TCGABiolinksGUI tool (Silva et al., 2018) to run ELMER via a web-based interface, (iv) output of all results in a single interactive HTML file include all data tables, figures, and source code, (v) adoption of software engineering best practices including unit testing and better exception handling, (vi) annotation of cell-type specific chromatin context for resulting genomic elements, and (vii) a new *Supervised* mode where the user can explicitly define sample groups for comparison. In this brief Note, we highlight several of these new features by analyzing TCGA Breast Cancer data to identify molecular subtype-specific networks. A complete description of new methods and features, along with computational benchmarking, is presented in the Supplementary Methods and Notes (Supplementary Figures 1–16 and Supplementary Tables S1–S5). ELMER v. 2 has been publicly available starting with v. 2.2.7 in Bioconductor Release 3.6 (October 2017). Complete result reports for the BRCA analyses are available in the Supplemental Materials and at http://bit.ly/ELMER_reports.

**Figure 1.**
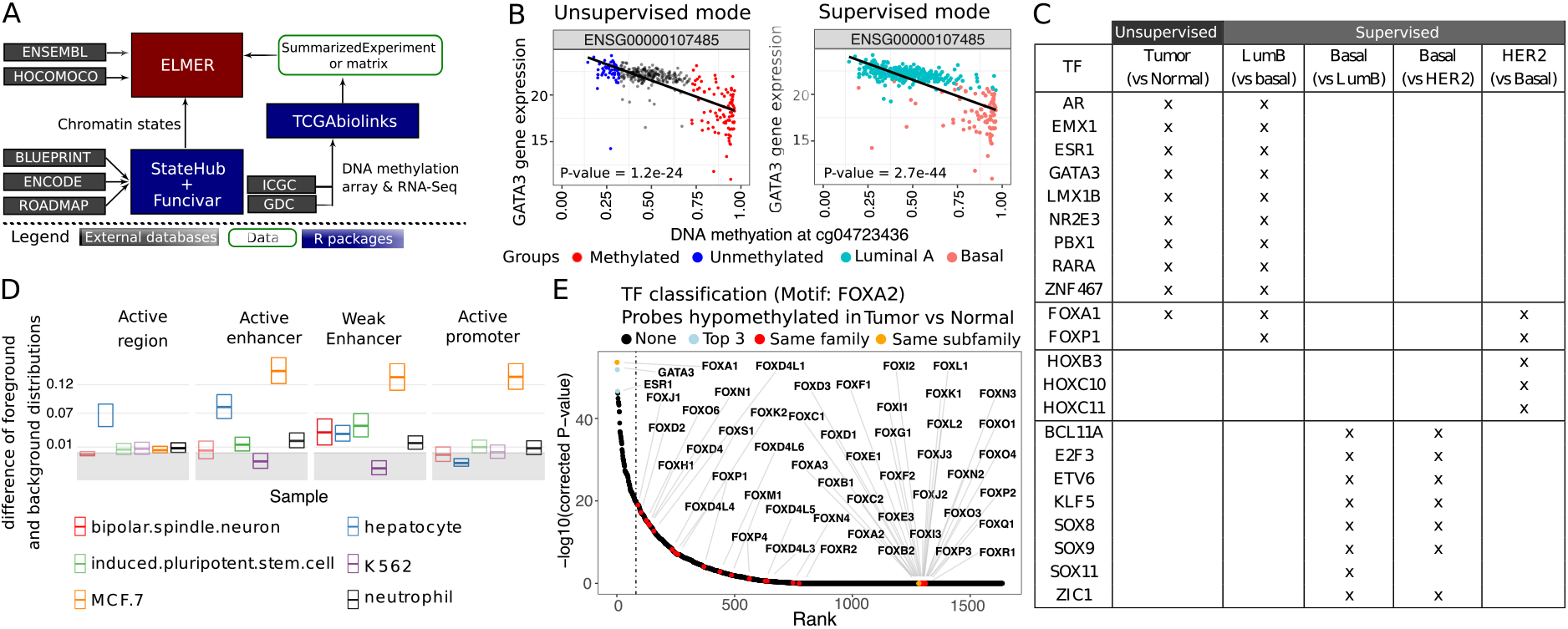
(A) ELMER architecture, showing external data sources (gray) and Bioconductor packages (blue). (B) Association of enhancer probe methylation and expression of the nearby *GATA3* gene, showing sample groups used in the *Unsupervised* vs. *Supervised* analysis modes. In *Unsupervised* mode, the 20% of samples with the lowest (blue) and highest (red) methylation levels are compared; in *Supervised* mode, the predefined Luminal A (blue) and Basal-like (red) tumors are compared. (C) A selected set of subtype-specific Master Regulator candidates identified from TCGA BRCA, comparing *Unsupervised* vs. various *Supervised* analysis runs. The complete table is available as Supplementary Table S3. (D) StateHub chromatin state enrichment analysis for 1, 076 regulatory elements identified in the *Unsupervised* analysis. (E) Master Regulator analysis for the top motif in the *Unsupervised* analysis, *FOXA2*. All TFs are ranked by their correlation with methylation changes of distal probes within 250 bp of a *FOXA2* binding motif. Colored dots indicate the top 3 most anti-correlated TFs (*FOXA1, GATA3* and *ESR1*), and all TFs classified in the same family as *FOXA2*.

## 2 Feature highlights

### Supervised vs Unsupervised mode

ELMER first identifies Differentially Methylated CpGs (DMCs) occurring at distal (non-promoter) probes (Step 1), then searches for downstream gene targets for each DMC (Step 2), and finally identifies Master Regulator TFs based on enriched binding motifs and TF expression (Step 3), as shown in Supplementary Fig. 1. ELMER v. 1 identified DMCs by comparing methylation in all cancer vs. non-cancer samples, while the subsequent steps used correlation between methylation and expression in the *n*% of tumors with the most extreme methylation values (by default, *n*=20). The rationale was that any particular GRN might only be altered in a subset of tumors with a specific molecular phenotype, which would not always be known *a priori*. While 20% was an arbitrary definition, we found this to be a useful exploratory strategy given the heterogeneity of cancer molecular phenotypes.

In ELMER v. 2, we continue to support this original *Unsupervised* strategy. However, we have found many practical use cases where the group structure is known in advance, and a *Supervised* search strategy is preferable. This is especially true for “case-control” experimental designs such as treated vs. untreated samples. The major difference is that in *Supervised* mode, all samples must be contained in one of the two comparison groups, whereas *Unsupervised* mode still uses only the *n*% most extreme. Furthermore, this subset of samples with the most extreme methylation values changes from one genomic locus to the next.

To compare *Supervised* vs. *Unsupervised* modes, we used ELMER v. 2.4.3 to analyze TCGA BRCA (Breast Invasive Carcinoma) data (Supplementary Figures 2–15 and Supplementary Tables 2–3). Based on enhancer-gene pairing, *Unsupervised* mode had lower statistical power (Fig. 1B), but was able to identify molecular subtype-specific networks without explicit *a priori* subtype labels (Fig. 1C). As expected, *Supervised* mode is best suited to explore well-understood molecular phenotypes, while *Unsupervised* mode can be a powerful tool to discover networks in unknown tumor subtypes. When molecular subtypes are known, the two modes can be used in conjunction and compared (as we have done in Supplemental Table S3).

### Functional interpretation of chromatin states

While ELMER v.1 was limited to analyze only probes overlapping known enhancers, ELMER v.2 analyzes *all* distal probes, and thus it is now important to provide a functional interpretation of the resulting regions. We perform a chromatin state enrichment analysis using states automatically downloaded from the (http://StateHub.org) database, a publicly-available resource that integrates histone modification and other publicly-available epigenomic data for over 1,000 different human samples (Coetzee et al., 2018). Enrichment of these states is calculated against a randomly sampled background set drawn from the same distal probe set used as input. We used ELMER 2 to perform this state enrichment analysis for the BRCA dataset, yielding insights into the cell-type specificity of the genomic regions identified (Fig. 1D, and Supplementary Fig. 5). The strongest enrichment was for active enhancer and promoter states having cell-type specificity for MCF7, a Luminal Breast Cancer cell line.

### Motif enrichment analysis and identification of Master Regulator TFs

The final step of ELMER identifies enriched TF binding motifs within candidate regulatory regions, followed by correlation with TF expression to identify upstream Master Regulators (Supplementary Fig. 1). ELMER v. 1 used a hand-curated selection of 145 TF motifs, which were grouped into binding domain families manually. We re-implemented these sections in ELMER v. 2 to use publicly available databases for these steps, making the package more comprehensive and easier to update in future versions. ELMER v. 2 uses 771 human binding models from HOCOMOCO v11 (Kulakovskiy et al., 2017). Each of these is associated with one or more of 1,639 transcription factors defined in (Lambert et al., 2018), which are grouped into 82 different binding domain families and 331 sub-families using the TFClass database (Wingender et al., 2017). We use the Fisher’s exact test and Benjamini-Hochberg multiple hypothesis correction to compare the frequency of each motif flanking the positive CpG probes to a background defined by all distal probes on the array, plotting the top hits as odds ratios with 95% confidence intervals (Supplementary Fig. 13).

For each enriched motif, we then calculate a mean DNA methylation value for all probes having a motif instance within ±250*bp*, and correlate this value to each of the 1, 639 TFs in our database. This helps to distinguish between different members of the same TF family, which often have nearly indistinguishable binding motifs. For instance, in the BRCA analysis, the most highly enriched motif corresponded to *FOXA2*, but our this Master Regulator (MR) analysis showed the likely family member to be *FOXA1* (Fig. 1E), which has been extensively validated as a MR in luminal subtypes of breast cancer (Meyer and Carroll, 2012; Nakshatri and Badve, 2009). We ran the same analysis with the *Supervised* mode to compare explicit changes in each of the known molecular subtypes from (Ciriello et al., 2015), which had a significant overlap with the *Unsupervised* analysis but yielded many additional MRs (Fig. 1C, Supplementary Table S3). Two examples of were *SOX11* and *KLF5*, whose functional roles in basal-like BRCA were recently described (Shepherd et al., 2016; Ben-Porath et al., 2008), and Androgen Receptor (*AR*), which has been implicated in ER-positive BRCA (Feng et al., 2017; Vera-Badillo et al., 2013). In addition to these known regulators, many completely unexplored TFs were identified as candidate MRs (Supplementary Table S3), highlighting the power of *Unsupervised* analysis.

## 3 Conclusions and Future Directions

ELMER v. 2 has been substantially re-written based on Bioconductor standards and user needs. The new *Supervised* mode and improved TF analysis identified additional known and novel Master Regulators candidates in TCGA BRCA analyses. ELMER v. 2 has only been tested on data from Illumina methylation arrays, which cover only 5-15% of all enhancer regions based on whole-genome bisulfite sequencing (WGBS). While *ELMER* does not currently support WGBS due to lack of sufficient test data, the number of WGBS datasets is quickly growing, and we expect the same basic ELMER approach will scale well in the future to take advantage of this more comprehensive data type.

## Introduction

In addition to the details below, a complete HTML output report for the two runs described in the *Use case* Section is available at http://bit.ly/ELMERv2_reports. This document contains all source code, parameters used, Methods descriptions, output tables, and output plots.

### ELMER workflow

The complete ELMER workflow is shown in Supplementary Fig. 1.

**Supplementary Fig. 1.**
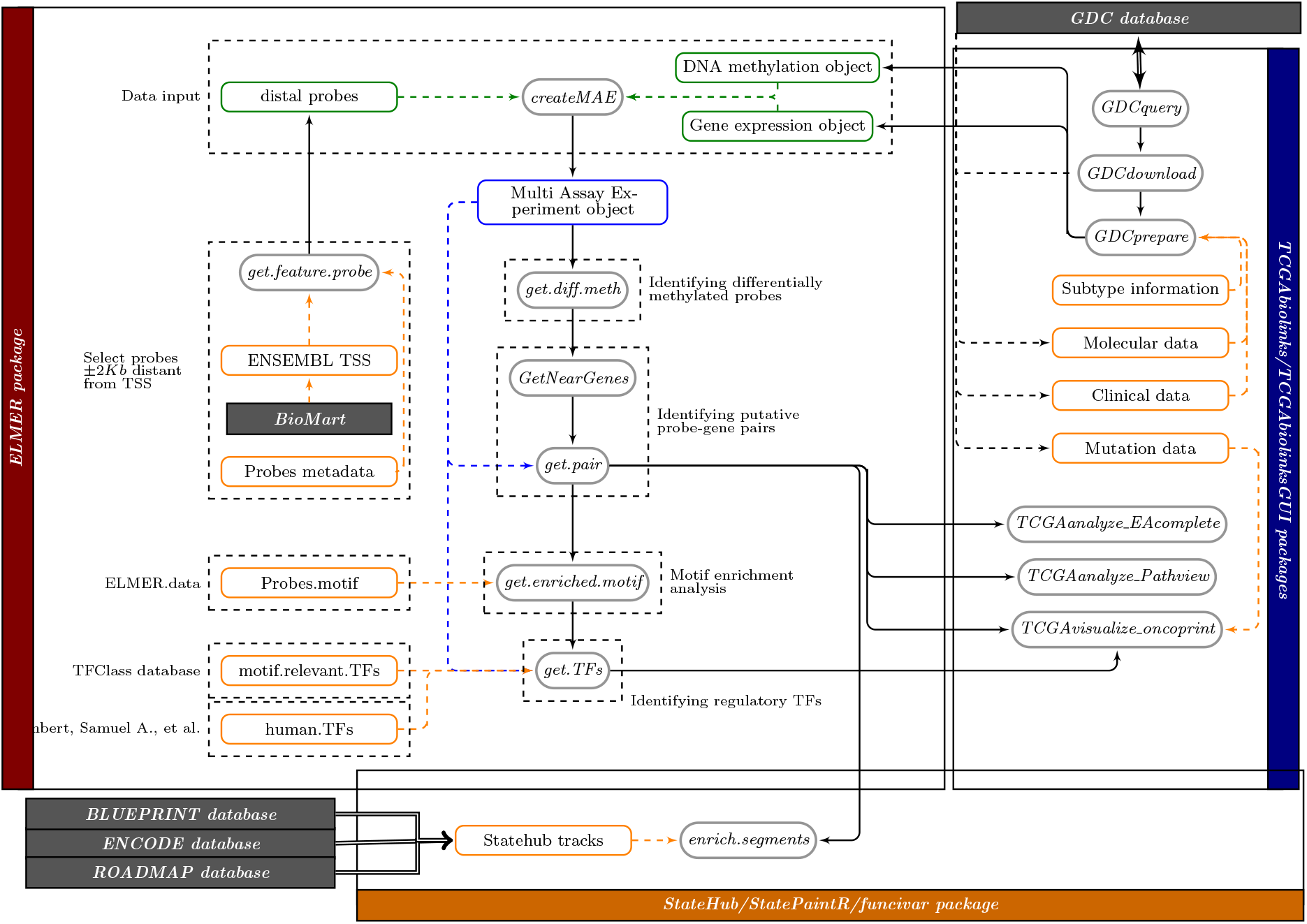
ELMER workflow: ELMER receives as input a DNA methylation array object and a gene expression object (matrices or SummarizedExperiment objects) and a Genomic Ranges (GRanges) object with distal probes to be used as filter which can be retrieved using the *get.feature.probe* function. The function *createMAE* will create a Multi Assay Experiment object keeping only samples that have both DNA methylation and gene expression data. Genes will be mapped to genomic position and annotated using ENSEMBL database (Aken et al., 2016), while for probes it will add annotation from Zhou et al. (http://zwdzwd.github.io/InfiniumAnnotation). This MAE object will be used as input to the next analysis functions. First, it identifies differentially methylated probes followed by the identification of their nearest genes (10 upstream and 10 downstream) through the *get.diff.meth* and *GetNearGen.es* functions respectively. For each probe, it will verify if any of the nearby genes were affected by its change in the DNA methylation level and a list of gene and probes pairs will be outputted from *get.pair* function. For the probes in those pairs, it will search for enriched regulatory Transcription Factors motifs with the *get.enriched.motif* function. Finally, the enriched motifs will be correlated with the level of the transcription factor through the *get. TFs* function. In the figure green Boxes represents user input data, blue boxes represent output object, orange boxes represent auxiliary pre-computed data and gray boxes are functions.

### Main differences between ELMER old version (ELMER 1) and the new version (ELMER 2)

The main differences between ELMER 1 and ELMER 2 are summarized in the Supplementary Table S1.

**Table S1.** Main differences between ELMER old version (v.1) and the new version (v.2)

| Features | ELMER Version 1 | ELMER Version 2 |
| --- | --- | --- |
| Primary data structure | mee object (custom data structure) | MAE object (Bioconductor data structure) |
| Auxiliary data | Manually created | Programmatically created |
| Number of human TFs | 1,982 | 1,639 (curated list from Lambert, Samuel A., et al.) |
| Number of TF motifs | 145 | 771 (HOCOMOCO v11 database) |
| TF classification | 78 families | 82 families and 331 subfamilies (TFClass database, HOCOMOCO) |
| Analysis performed | Normal vs tumor samples | Group 1 vs group 2 |
| Statistical grouping | Unsupervised only | Unsupervised or supervised using labeled groups |
| TCGA data source | The Cancer Genome Atlas (TCGA) (not available) | The NCI's Genomic Data Commons (GDC) |
| Genome of reference | GRCh37 (hg19) | GRCh37 (hg19)/GRCh38 (hg38) |
| DNA methylation platforms | HM450 | EPIC and HM450 |
| Graphical User Interface (GUI) | None | TCGAbiolinksGUI |
| Automatic report | None | HTML summarizing results |
| Annotations | None | StateHub |

### Organization of data as a *MultiAssayExperiment* object

To facilitate the analysis of experiments and studies with multiple samples, the Bioconductor team created the *SummarizedExperiment* class (Huber et al., 2015), a data structure able to store data and metadata for a single experiment but not for data spanning several experiments for the same sample. To overcome this problem, recently, the MultiAssay Special Interest Group (SIG) created the *MultiAssayExperiment* class (Ramos et al., 2017) a data structure to manage and preprocess multiple assays for integrated genomic analysis. This data structure is now an input for all main functions of ELMER and can be generated by the *createMAE* function.

To perform *ELMER* analyses, we populate a *MultiAssayExperiment* with a DNA methylation matrix or *SummarizedExperiment* object from HM450K or EPIC platform; a gene expression matrix or SummarizedExperiment object for the same samples; a matrix mapping DNA methylation samples to gene expression samples; and a matrix with sample metadata (i.e. clinical data, molecular subtype, etc.). TCGA or other GDC data can be imported by TCGAbiolinks (Supplementary Fig. 1), in which case the necessary data structures are automatically created. Based on the genome of samples selected, metadata for the DNA methylation probes, such as genomic coordinates, are added from (Zhou et al., 2016); and metadata for gene annotation is added from the ENSEMBL database (Yates et al., 2015) using biomaRt (Durinck et al., 2009). Use of these standardized import packages allows ELMER v.2 to take advantage of all current datasets. For instance, TCGABiolinks will soon be able to read from the International Cancer Genome Consortium (ICGC) repository, and similar importers can be written for other disease databanks.

If using non-TCGA data, the matrix with sample metadata should be provided with at least a column with a subject identifier and another one identifying its group which are used for analysis, if samples in the methylation and expression matrices are not ordered and with same names, a matrix mapping for each patient identifier their DNA methylation samples and their gene expression samples should be provided to the *createMAE* function.

### Selecting distal probes

Probes from HumanMethylationEPIC (EPIC) array and Infinium HumanMethylation450 (HM450) array are removed based on the default filtering manifests from (Zhou et al., 2017). Briefly, probes are masked from the analysis if they have either internal SNPs close to the 3′ end of the probe; non-unique mapping to the bisulfite-converted genome; or off-target hybridization due to partial overlap with non-unique elements. Probe metadata information is available as ELMER.data package, populated from the source file at http://zwdzwd.github.io/InfiniumAnnotation (Zhou et al., 2017).

For analysis of distal elements, probes located in regions of ±2*kb* around transcription start sites (TSSs) are removed.

### Supervised vs. Unsupervised modes

ELMER is designed to identify differences between two sets of samples within a given dataset. In Yao et al., the first step (identification of DMCs) was hard-coded to identify DMCs between non-cancer vs. cancer samples, and the subsequent step was *unsupervised*, identifying changes within any subset of tumors. In ELMER v.2, we generalize these strategies so that they are applicable to any paired dataset, including disease vs. healthy tissue for any disease type, untreated vs. treated samples, etc. We now support two modes, with the *Unsupervised* mode based on the original method from Yao et al.. Here, the user defines Group 1 and Group 2 samples, but an assumption is made that only a subset of samples differs between the two groups. By default, this subset includes the most extreme 20% of samples within the group, and this is an input parameter can be modified. The new mode is the *Supervised* mode, in which *all* available samples from each group are used. This mode should be used when pre-determined phenotypes or molecular subtypes are known in advance, such as the treated vs. untreated case. The advantage is that this greatly increases statistical power because of all samples from each group. This can be extremely important, given the large burden of multiple hypothesis testing involved in ELMER.

### Identification of differentially methylated CpGs (DMCs)

The first step is the identification of differentially methylated CpGs (DMCs). In the *Supervised* mode, we compare the DNA methylation level of each distal CpG for *all* samples in Group 1 compared to all samples Group 2, using an unpaired one-tailed t-test. In the *Unsupervised* mode, the samples of each group (Group 1 and Group 2) are ranked by their DNA methylation beta values for the given probe, and those samples in the lower quintile (20% samples with the lowest methylation levels) of each group are used to identify if the probe is hypomethylated in Group 1 compared to Group 2. The reverse applies for the identification of hypermethylated probes. It is important to highlight that in the *Unsupervised* mode, each probe selected may be based on a different subset the samples, and thus probe sets from multiple molecular subtypes may be represented. In the *Supervised* mode, all tests are based on the same sample grouping.

The 20% is a parameter to the *diff.meth* function called *minSubgroupFrac*. For the unsupervised analysis, this is set to 20% as in (Yao et al., 2015), because we wanted to be able to detect a specific molecular subtype among samples; these subtypes often make up only a minority of samples, and 20% was chosen as a lower bound for the purposes of statistical power (high enough sample numbers to yield t-test p-values that could overcome multiple hypotheses corrections, yet low enough to be able to capture changes in individual molecular subtypes occurring in 20% or more of the cases.) This number can be set as an input to the *diff.meth* function and should be tuned based on sample sizes in individual studies. The parameter value is always shown in the Settings section of the ELMER HTML output report. In the *Supervised* mode, where the comparison groups are implicit in the sample set and labeled, the *minSubgroupFrac* parameter is set to 100%. An example would be a cell culture experiment with 5 replicates of the untreated cell line, and another 5 replicates that include an experimental treatment.

To identify hypomethylated DMCs, a one-tailed t-test is used to rule out the null hypothesis: *μ_group1_* ≥ *μgroup*_2_, where *μ_group1_* is the mean methylation within the lowest group 1 quintile (or another percentile as specified by the *minSubgroupFrac* parameter) and *μ_group2_* is the mean within the lowest group 2 quintile. Raw p-values are adjusted for multiple hypothesis testing using the Benjamini-Hochberg method (Benjamini and Hochberg, 1995), and probes are selected when they had adjusted p-value less than 0.01 (which can be configured using the *pvalue* parameter). For additional stringency, probes are only selected if the methylation difference: Δ = *μ_group1_ − μ_group2_* was greater than 0.3. This can be configured with the *sig.diff* parameter. The same method is used to identify hypermethylated DMCs, except we use the *upper* quintile, and the opposite tail in the t-test is chosen.

### Identification of putative target gene(s)

For each differentially methylated distal probe (DMC), the closest 10 upstream genes and the closest 10 downstream genes are tested for inverse correlation between methylation of the probe and expression of the gene, which is the same basic strategy employed in ELMER version 1. However, we now import all gene annotations programmatically using the Biomart (Durinck et al., 2005, 2009) package. This allows easy extensibility to use any annotations desired (our default uses Ensembl annotations).

This step also differs between the *Supervised* and *Unsupervised* modes. In the *Unsupervised* mode, as in ELMER v.1, for each probe-gene pair, the samples (all samples from both groups) are divided into two groups: the M group, which consist of the upper methylation quintile (the 20%of samples with the highest methylation at the enhancer probe), and the U group, which consists of the lowest methylation quintile (the 20% of samples with the lowest methylation). In the new *Supervised* mode, the U and M groups are defined strictly by sample group labels, and all samples in each group are used. The *Supervised* mode can greatly increase statistical power, as illustrated in Supplementary Fig. 2.

For each differentially methylated distal probe (DMC), the closest 10 upstream genes and the closest 10 downstream genes are tested for inverse correlation between methylation of the probe and expression of the gene (the number 10 can be changed using the *numFlankingGenes* parameter). To select these genes, the probe-gene distance is defined as the distance from the probe to the transcription start site specified by the ENSEMBL gene level annotations (Yates et al., 2015) accessed via the R/Bioconductor package biomaRt (Durinck et al., 2009, 2005). By choosing a constant number of genes to test for each probe, our goal is to avoid systematic false positives for probes in gene rich regions. This is especially important given the highly non-uniform gene density of mammalian genomes.

Thus, exactly 20 statistical tests were performed for each probe, as follows.

For each candidate probe-gene pair, the Mann-Whitney U test is used to test the null hypothesis that overall gene expression in group M is greater than or equal than that in group U. This non-parametric test was used in order to minimize the effects of expression outliers, which can occur across a very wide dynamic range.

In the *Unsupervised* mode, for each probe-gene pair tested, the raw p-value *P_r_* is corrected for multiple hypothesis using a permutation approach as follows. The gene in the pair is held constant, and *x* random methylation probes are chosen to perform the same one-tailed U test, generating a set of *x* permutation p-values *P_p_*. We chose the x random probes only from among those that were “distal” (farther than 2*kb* from an annotated transcription start site), in order to draw these null-model probes from the same set as the probe being tested (Sham and Purcell, 2014). An empirical p-value *P_e_* value was calculated using the following formula (which introduces a pseudo-count of 1):

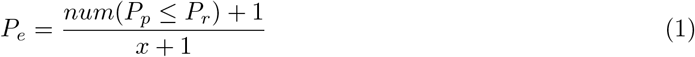

In the supervised mode, for each probe-gene pair tested, the raw p-value *P_r_* is corrected for multiple hypothesis using Benjamini-Hochberg procedure. Also, notice that in the *Supervised* mode, no additional filtering is necessary to ensure that the *M* and *U* group segregate by sample group labels. The two sample groups are segregated by definition, since these probes were selected for their differential methylation, with the same directionality, between the two groups.

**Supplementary Fig. 2.**
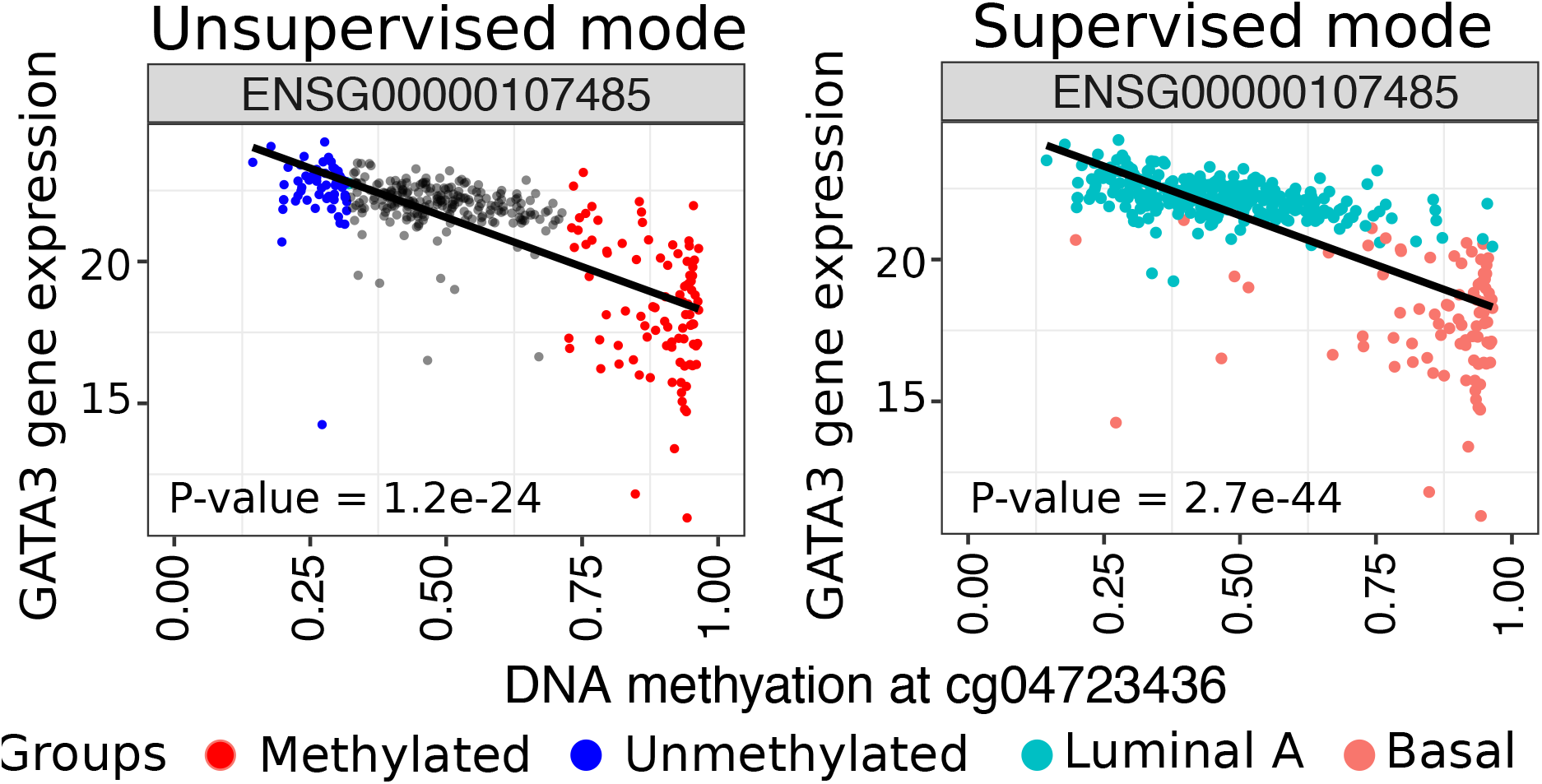
*Supervised* mode maximizes statistical power. Difference of groups U and M definition in *Supervised* and *Unsupervised* mode. A: *Unsupervised* mode; when *minSubgroupFrac* argument is set to 40%, the methylated group is defined as the highest quintile and the unmethylated group as the lowest quintile; B: *Supervised* mode; methylated and unmethylated group are defined as one of the known molecular subtypes. For example, the unmethylated group is represented by all the LumA samples while the methylated group is represented by all the Basal samples. The t-test p-value achieved for the Unsupervised mode is 1.2*E* − 24, while the Supervised mode is: 2.7*E* − 44.

### Characterization of chromatin state context of enriched probes using StateHub

While ELMER version 1 was limited to searching within annotated enhancer elements, we have since found that this constraint was not necessary to achieve statistical power. Thus ELMER v.2 by default searches *all* distal elements in the genome (distal elements are those greater than ±2*kb* from a TSS; By changing ELMER default settings, it is possible to analyze TSS-proximal probes, either together with distal probes or separately. See ELMER Bioconductor documentation for details).

Because ELMER can now search essentially all probes on the array, it is important to understand the context of the probes that result from an ELMER analysis. Typically, these are enhancer probes, but some regulatory changes may involve unannotated promoters, insulators, etc. We used the *StateHub* (http://statehub.org/) (Coetzee et al., 2017) and *FunciVar* (https://github.com/Simon-Coetzee/funcivar) Bioconductor packages to characterize enrichment of the various cell-type-specific chromatin states in the significant BRCA-hypomethylated probes.

The Statehub Focused Poised Promoter Model (Decision Matrix) (Supplementary Fig. 4) is used to define the chromatin state of a region based on several marks. For example, an ‘‘Active region” (AR) is defined as overlapping one of the two ‘‘active” marks (either H3K9/14ac or H3K27ac) but neither the canonical promoter mark (H3K4me3) or the canonical enhancer mark (H3K4me1). If it has one of these marks, it is characterized either as an ‘Active Enhancer” (EAR) or ‘Active Promoter” (PAR). Also, a “Weak Enhancer” (EWR) state, has the enhancer regulatory mark (H3K4me1) but not the active mark H3K27ac. Also, Supplementary Fig. 3 shows the Statehub tracks used in the enrichment analysis and Supplementary Fig. 5 shows the its results.

Importantly, the MCF-7 cell line, and ER-positive breast cancer cell line, is much more strongly enriched for all enhancer and promoter classes than other cell types. As more reference cell types become available, this analysis will be useful in characterizing tumor GRN changes that reflect particular cell types or co-opted developmental programs.

All methods are described here: https://www.simoncoetzee.com/bioc2017.html FunciVar by default calculates a likelihood based on the beta-binomial distribution, returning a 95% credible interval (optionally set by the “CI” argument) for the range of differences between the two populations of variants (i.e. foreground and background). Specifically, it calculates a distribution of true enrichment (as probability of overlap) for both sets of variants in the genomic features based on the observed number of overlaps:

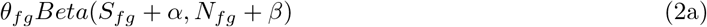

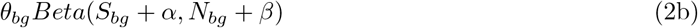

for S successes in N trials. FunciVar uses an uninformative Jeffreys prior c(*α*=0.5, *β*=0.5) to compare the two distributions directly by subtracting permuted samples to obtain the distribution of differences. The prior can be overridden in special cases.

**Supplementary Fig. 3.**
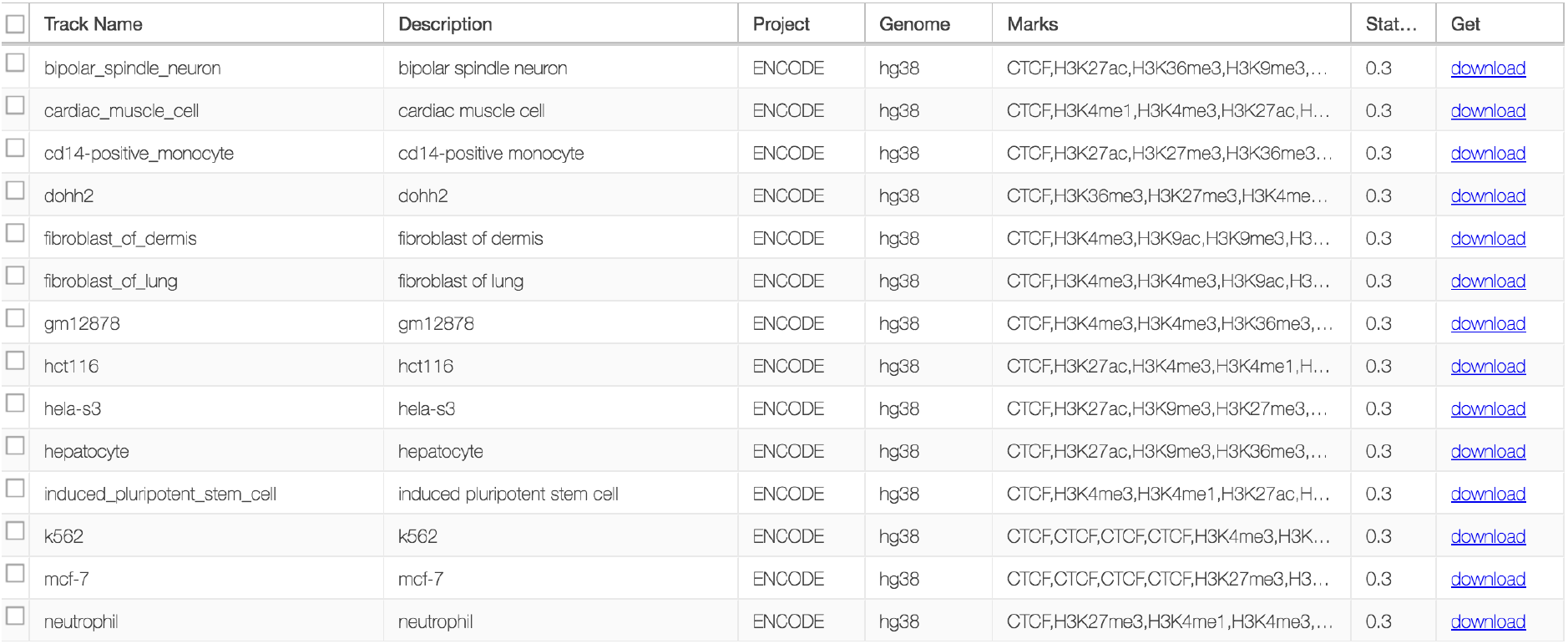
Statehub tracks for encode samples having H3K27ac, H3K4me1, H3K4me3 and CTCF marks for hg38 were used in Use cases. Retrieved from http://statehub.org/.

**Supplementary Fig. 4.**
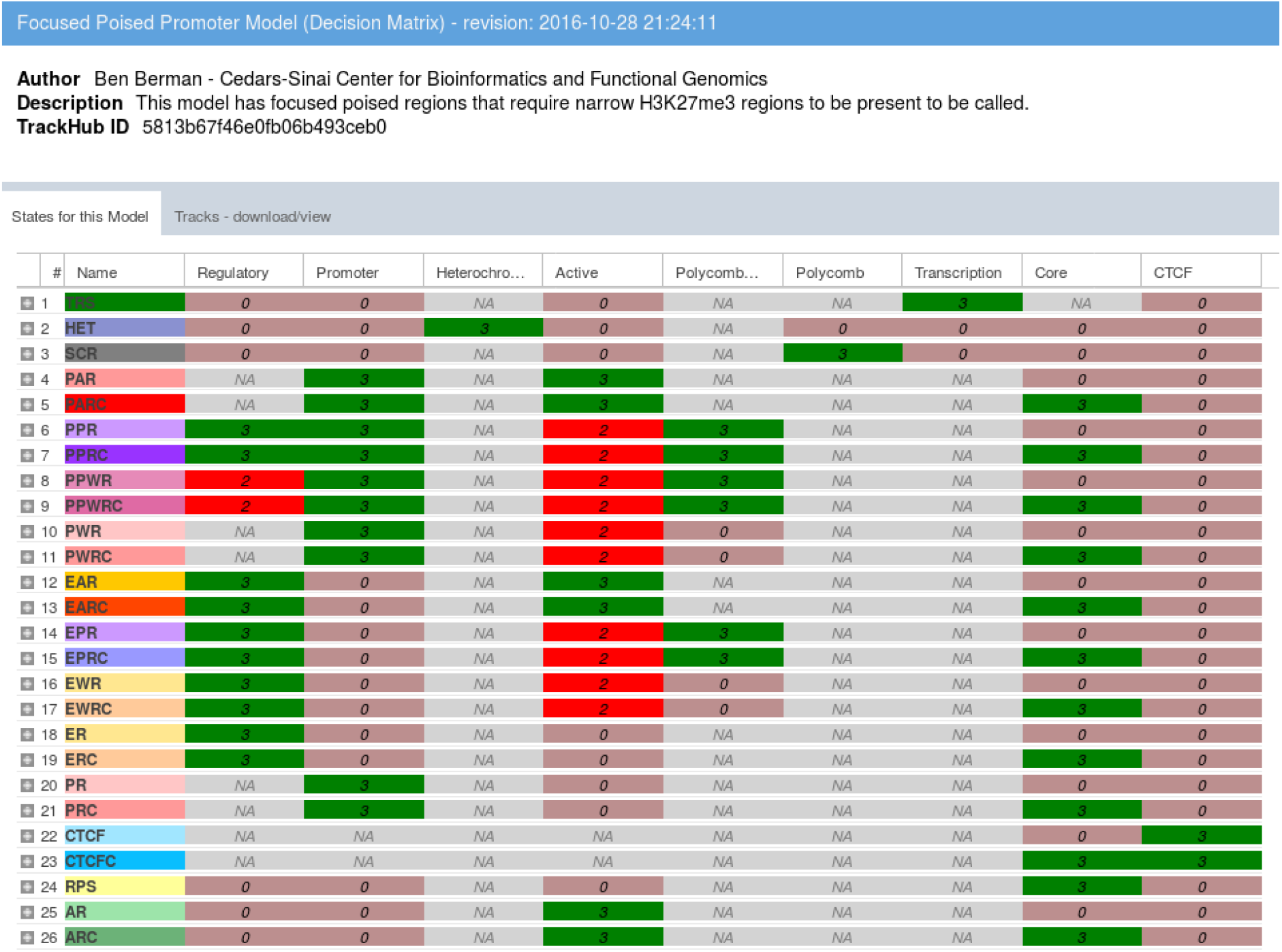
Tracks are available for approximately 1,000 human cell types in the StateHub portal, and some are shown in Supplemental Figure 3. States are defined by multi-mark model, such as the Focused Poised Promoter Model shown in Supplementary Figure 4. For example, an “Active region” (AR) is defined as overlapping one of the two “active” marks (either H3K9/14ac or H3K27ac) but neither the canonical promoter mark (H3K4me3) or the canonical enhancer mark (H3K4me1). If it has only one of these marks, it is characterized either as an “Active Enhancer” (EAR) or “Active Promoter” (PAR). Also, a “Weak Enhancer” (EWR) state, has the enhancer regulatory mark (H3K4me1) but not the active mark H3K27ac. Supplementary Figure 5 shows the enrichment plot.

**Supplementary Fig. 5.**
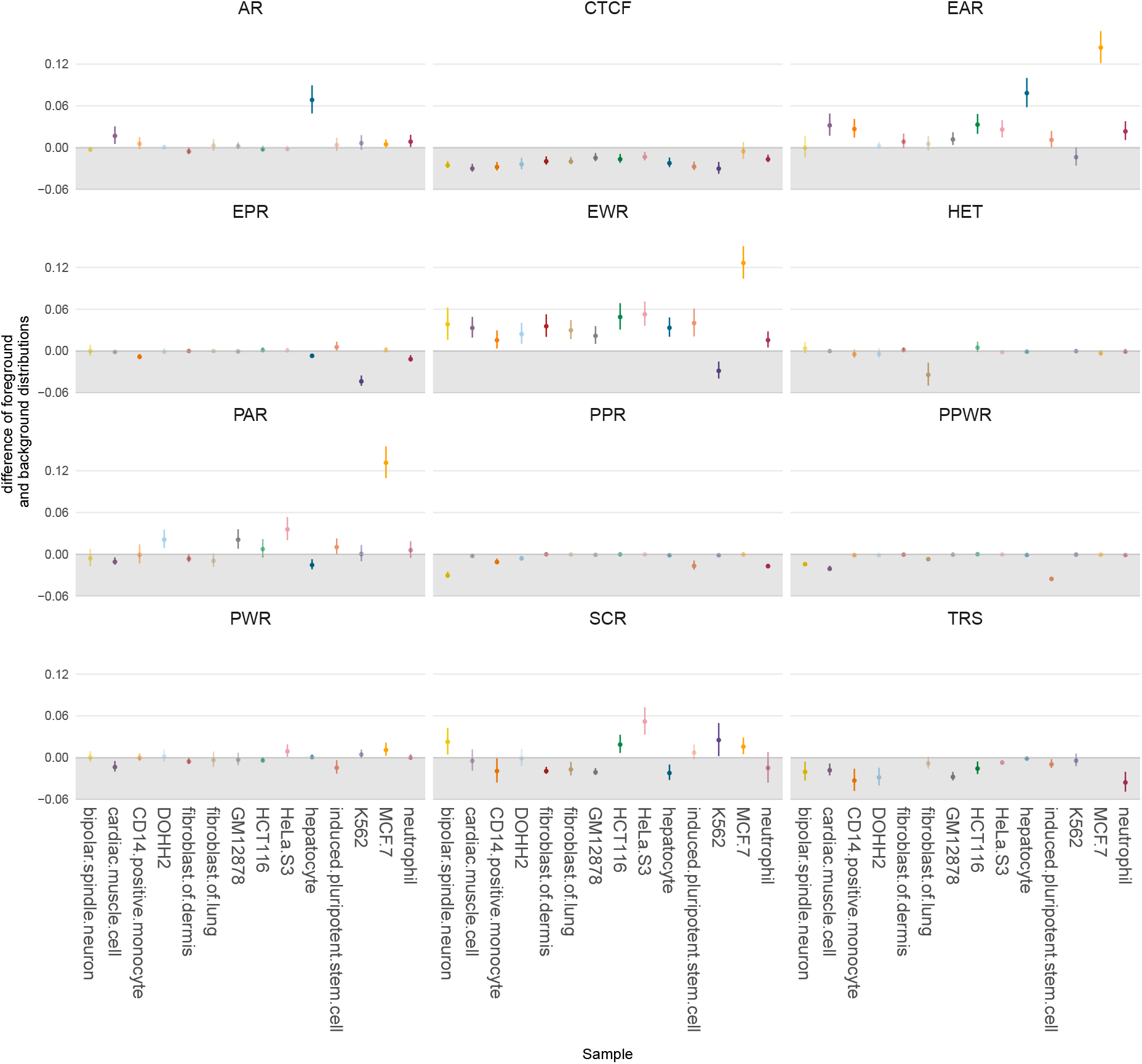
Enrichment of paired probes and chromatin states of ENCODE cells. The plot shows enrichment for enhancer active region, weak enhancer and active promoter region for MCF-7 cell. Acronyms - AR: Active region, EAR: active enhancer, EWR: Weak Enhancer, EPR: poised enhancer, PAR: active promoter, PWR: Weak Promoter, PPR: poised promoter, PPWR: Weak Poised Promoter, CTCF: architectural complex, TRS: transcribed, HET: heterochromatin, SCR: Polycomb Repressed Silenced. Y-axis shows the probability difference in overlap for the foreground class vs. random probes (Confidence Interval based on beta-binomial distribution, see methods).

### Motif enrichment analysis

In order to identify enriched motifs and potential upstream regulatory TFs, first, HOCOMOCO (HOmo sapiens COmprehensive MOdel COllection) v11 (Kulakovskiy et al., 2016, 2017) TF binding models, available at http://hocomoco.autosome.ru/downloads, were used as input for HOMER (Hypergeometric Optimization of Motif EnRichment) (Heinz et al., 2010) to find motif occurrences in a ±250*bp* region around each probe from EPIC and HM450 arrays.

For each probe set tested (i.e. the set of all probes occurring in significant probe-gene pairs), we quantify enrichments using Fisher’s exact test (where *a* is the number of probes within the selected probe set that contains one or more motif occurrences; *b* is the number of probes within the selected probe set that do not contain a motif occurrence; *c* and *d* are the same counts within the entire array probe set drawn from the same set of distal-only probes using the same definition as the primary analysis). Multiple testing correction with the Benjamini-Hochberg procedure (Fisher, 1922) is then applied to the Fisher’s results.

A probe set was considered significantly enriched for a particular motif if the 95% confidence interval of the Odds Ratio was greater than 1.1 (specified by option *lower.OR*, 1.1 is default), the motif occurred at least 10 times (specified by option *min.incidence*, 10 is default) in the probe set and *FDR* < 0.05.

### Identification of Master Regulator TFs

When a group of enhancers is coordinately altered in a specific sample subset, this is often the result of an altered upstream *master regulator* transcription factor in the gene regulatory network. *ELMER* identifies master regulator TFs corresponding to each of the TF binding motifs enriched from the previous analysis step. For each enriched motif, *ELMER* takes the mean DNA methylation of all distal probes (in significant probe-gene pairs) that contain that motif occurrence (within a ±250*bp* region), and compares this mean DNA methylation to the expression of each gene annotated as a human TF by (Lambert et al., 2018). The TFClass database (Wingender et al., 2013, 2017) is used to identify significantly associated TFs which are in the same DNA binding domain family or sub-family as the motif TF, information that is displayed in all output plots (Figure 1E) and HTML reports.

In the *Unsupervised* mode, a statistical test is performed for each motif-TF pair, as follows. All samples are divided into two groups: the M group, which consists of the 20% of samples with the highest average methylation at all motif-adjacent probes, and the *U* group, which consisted of the 20% of samples with the lowest methylation. This step is performed by the *get.TFs* function, which takes *minSubgroupFrac* as an input parameter, again with a default of 20%. For each candidate motif-TF pair, the Mann-Whitney U test is used to test the null hypothesis that overall gene expression in group *M* is greater or equal than that in group *U*. This non-parametric test was used in order to minimize the effects of expression outliers, which can occur across a very wide dynamic range. For each motif tested, this results in a raw p-value (*P_r_*) for each of the human TFs.

The new *Supervised* mode uses the same approach as described for the identification of putative target gene(s) step. The *U* and *M* groups are one of the the label group of samples and the *minSubgroupFrac* parameter is set to 100% to use all samples from both groups in the statistical test. This also can result in greater statistical power when using the *Supervised* mode.

### Comparing inferred results with MCF-7 ChIA-PET

As in our earlier paper (Yao et al., 2015), we compared CRM / gene pairs identified by Unsupervised analysis of TCGA Breast Cancer cases to chromatin loops derived from deep-sequenced ChIA-PET data from ER+ Breast Cancer MCF7 cells (Li et al., 2012). ELMER pairs were enriched for ChIA-PET loops by roughly 3-fold over random pairs (Supplementary Fig. 6), consistent with our earlier results.

First, we identify the number of ELMER pairs overlapping the ChIA-PET loops, then we repeat using randomly generated pairs with properties similar to the ELMER pairs. For each true ELMER probe in a probe-gene pair, we randomly select a different probe from the complete set of distal probes. We then choose the nth nearest gene to the random probe, where n is the same as the adjacency of the true ELMER probe (i.e. if the true probe is linked to the second gene upstream, the random probe will also be linked to its second gene upstream). Thus, the random linkage set has both the same number of probes and the same number of linked genes as the true set. One hundred such random datasets were generated to arrive at a 95% CI (±1.96 * *SD*). The result is shown in Supplementary Fig. 6. Of the 2118 putative pairs identified in breast cancer tumors, 223 (≈ 10.75%) were also identified as loops in the MCF7 ChIA-PET data. This was a three-fold enrichment over randomized probe-gene pairs.

**Supplementary Fig. 6.**
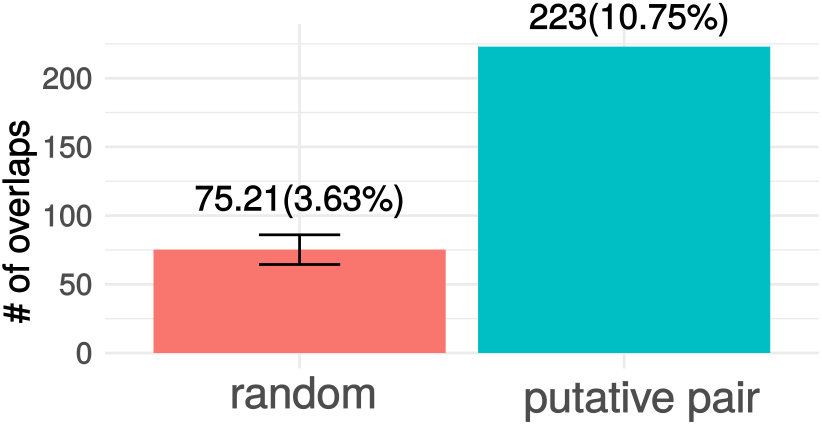
The graph shows the comparison of the number of probe-gene pairs identified within MCF7 ChIA-PET data using the putative pairs from BRCA vs. random pairs.

### Graphical User Interface

To enable user access to the methodologies offered in ELMER and to give users the flexibility of point-and-click style analysis without the need to learn R, we have implemented a full graphical user interface (GUI) through the R/Bioconductor package TCGAbiolinksGUI (Silva et al., 2017) available at http://bioconductor.org/packages/TCGAbiolinksGUI/. This tool allows definition of sample groupings based on user-defined clinical attributes in the supported databanks, including TCGA and TARGET, and GDC. A tutorial detailing the steps needed to use the tools through the GUI is available at https://bioinformaticsfmrp.github.io/Bioc2017.TCGAbiolinks.ELMER/index.html

ELMER can now be run directly from within TCGABiolinksGUI. Supplementary Figures 7,8 and 9 show the three ELMER menus.

**Supplementary Fig. 7.**
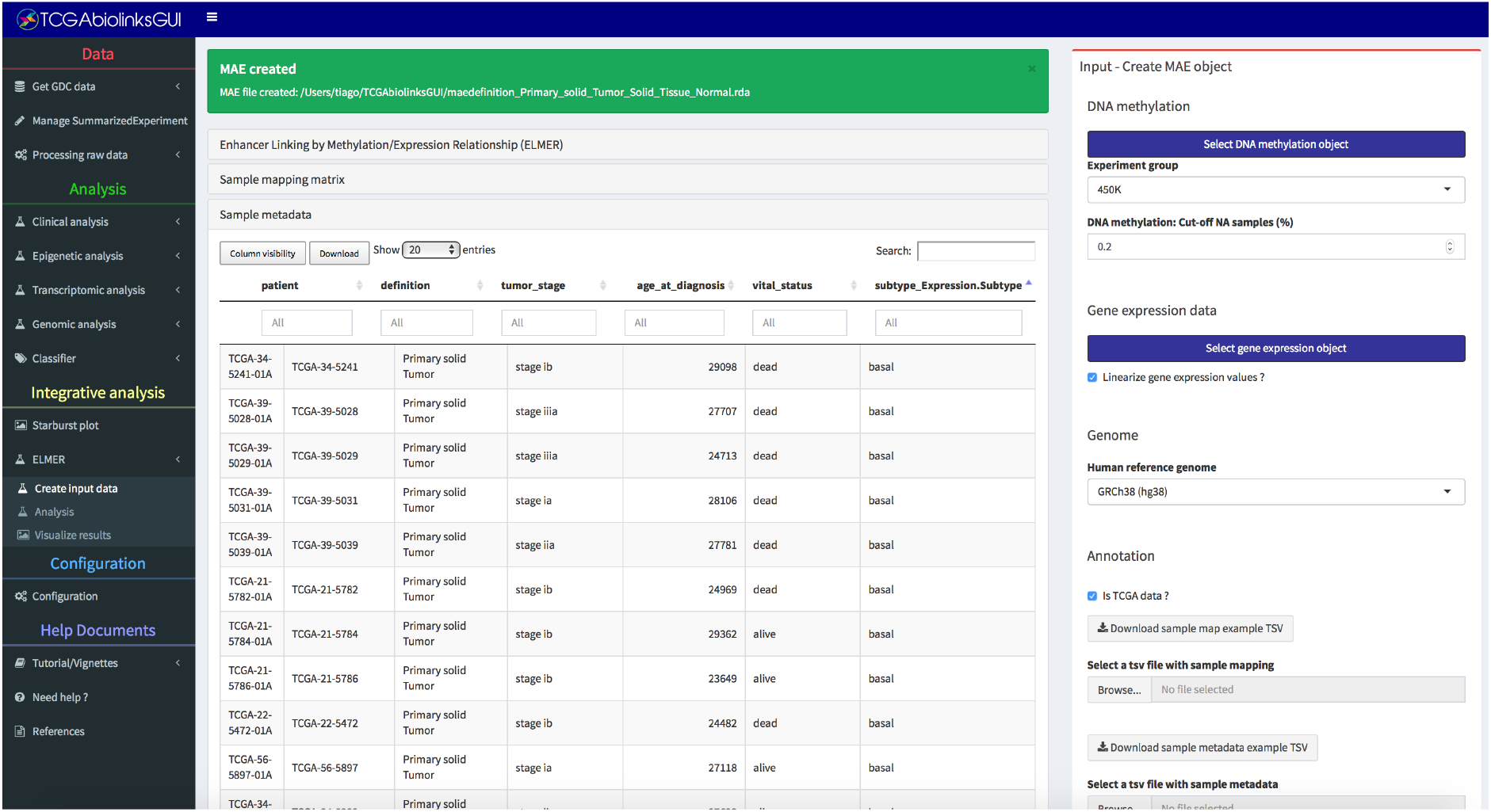
ELMER graphical user interface in TCGAbiolinksGUI: MAE creation menu.

**Supplementary Fig. 8.**
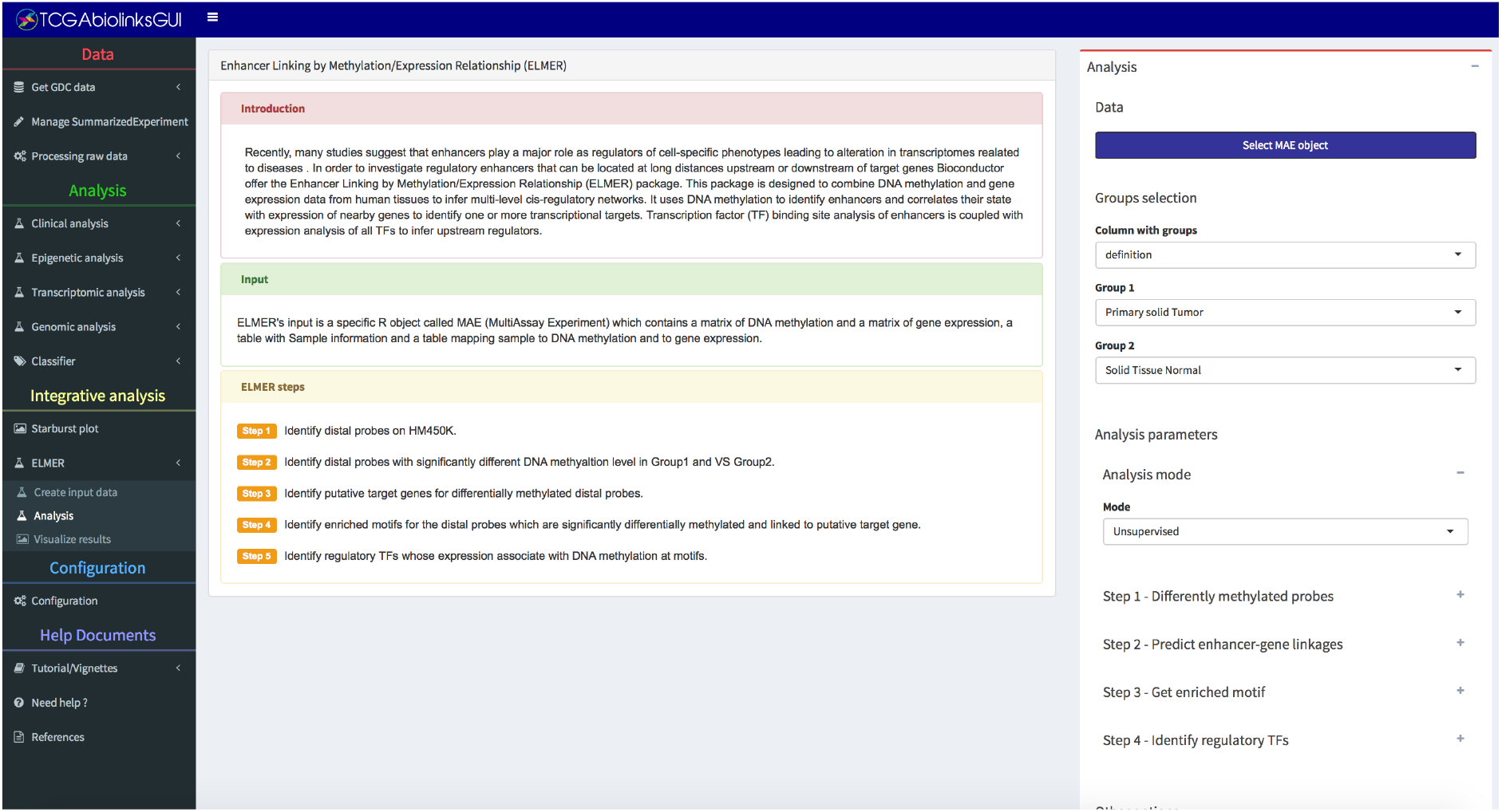
ELMER graphical user interface in TCGAbiolinksGUI: analysis menu.

**Supplementary Fig. 9.**
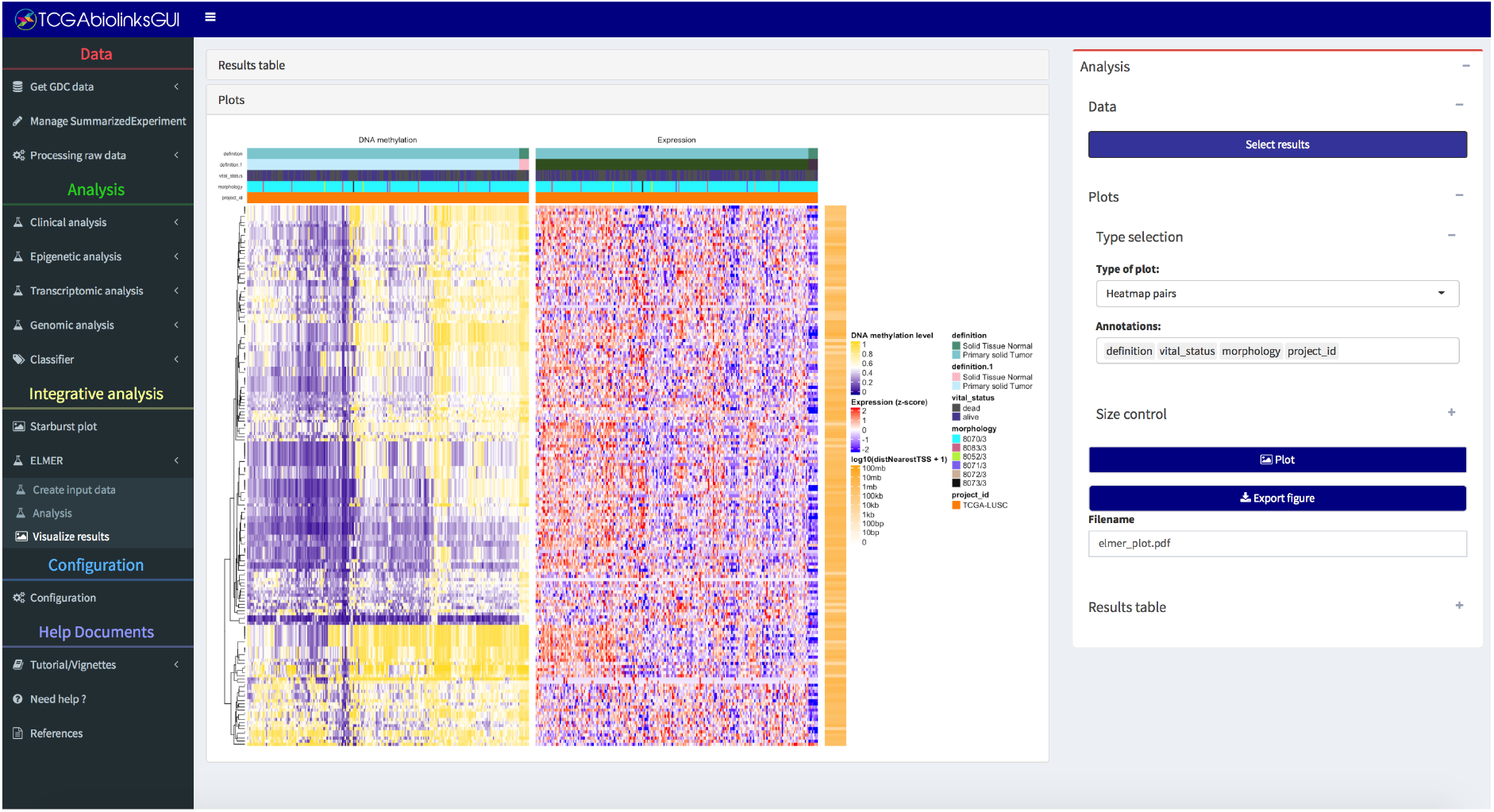
ELMER graphical user interface in TCGAbiolinksGUI: visualization menu.

### Interactive HTML output reports

While ELMER version 1 had functions to create individual output plots, we have completely revamped and added to the set of functions that create automatic output plots, which were used to generate all the Figures and Supplemental Figures in this article. We also now output a single HTML file which contains all source code used, output tables, and plots for an individual ELMER run. This HTML file is indexed via a table of contents, and individual sections can be expanded and compressed to expose additional detail.

Supplementary Figures 10 and 11 show the single interactive HTML file output for the TCGA Breast Cancer *Unsupervised* analysis described below. This and the HTML file for the Supervised analysis can both be downloaded at https://github.com/tiagochst/ELMER_supplemental/raw/master/supplemental_files.zip.

**Supplementary Fig. 10.**
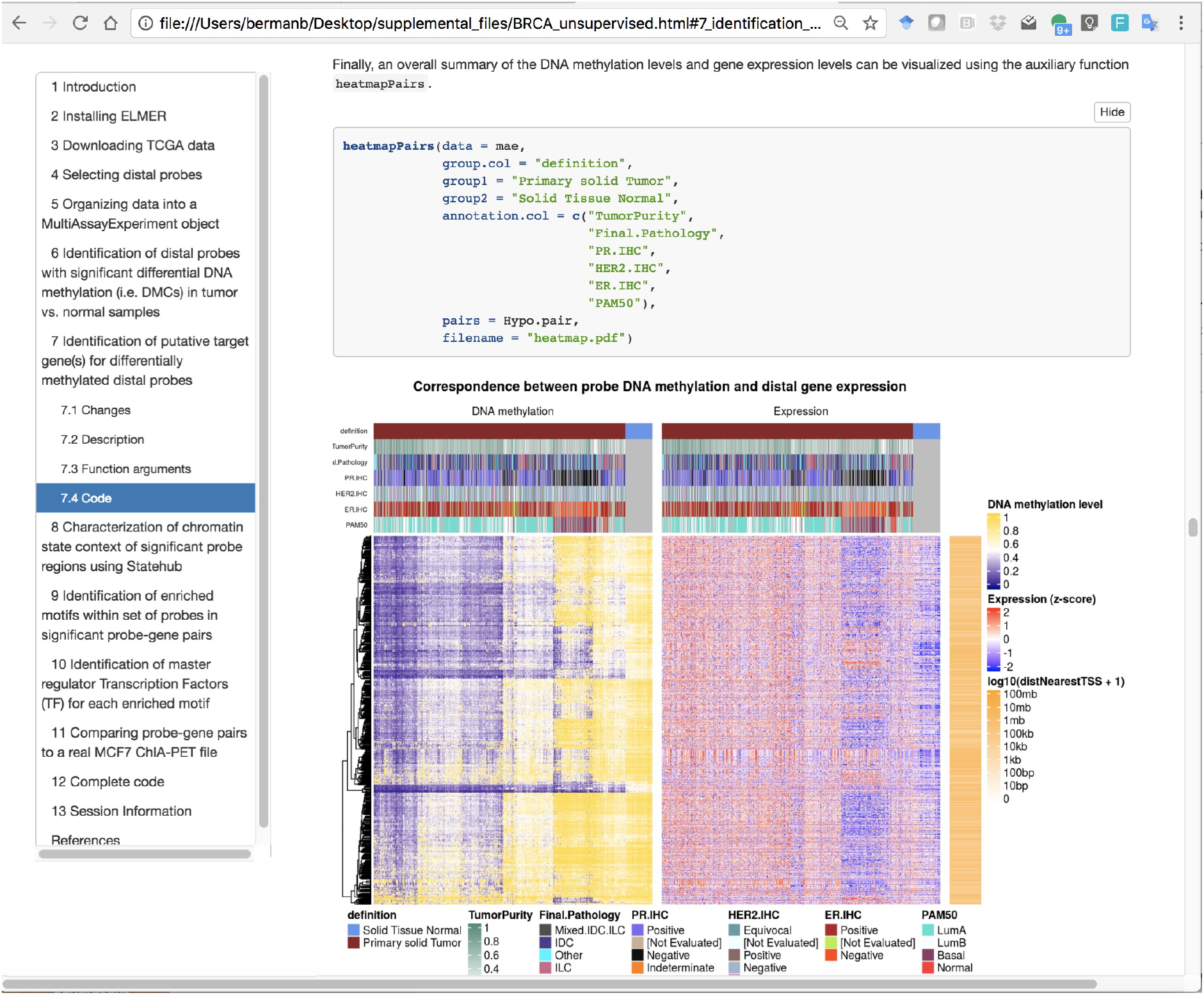
Single HTML file output report example, showing generation of the comprehensive heatmap plot.

**Supplementary Fig. 11.**
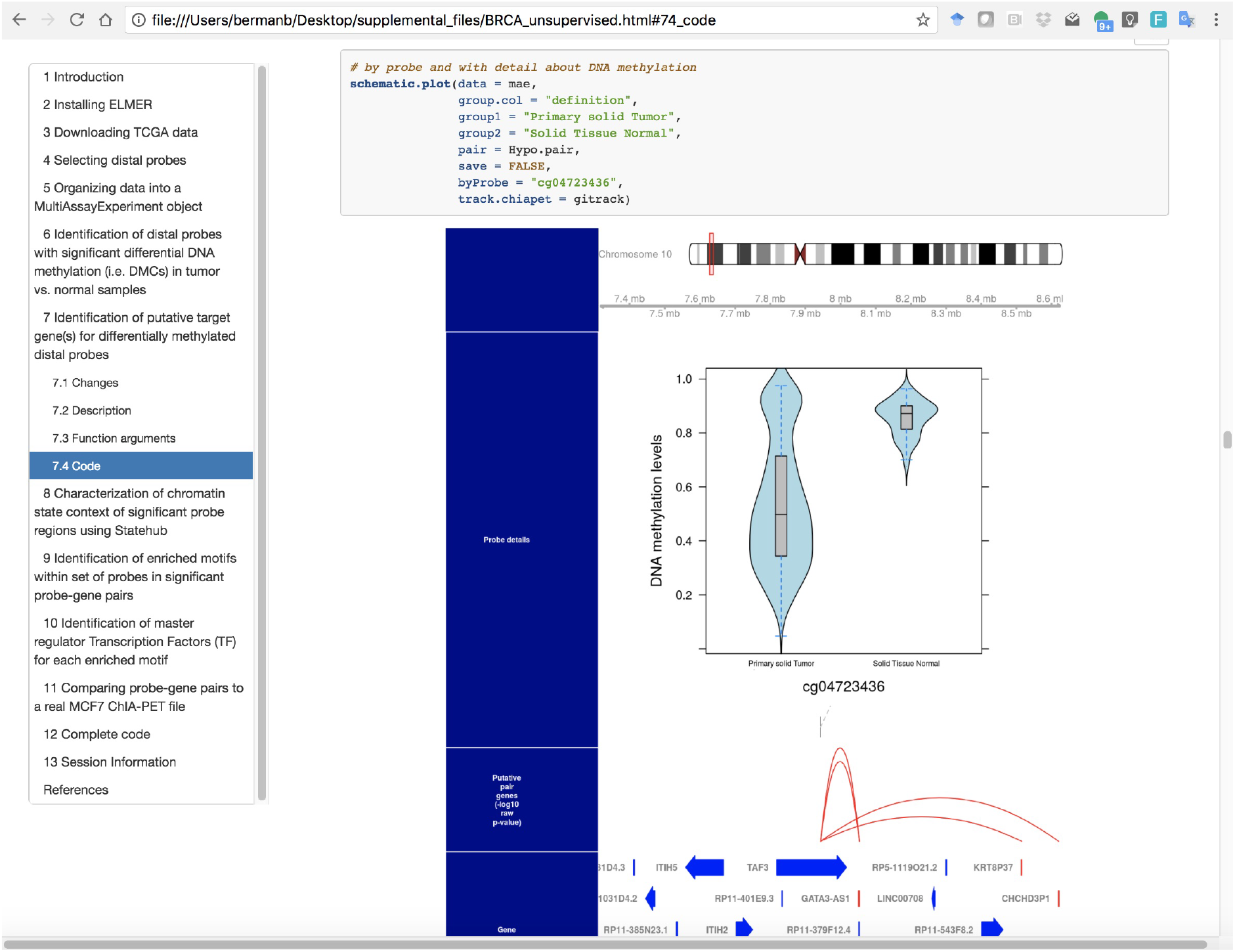
Single HTML file output report example, showing generation of the genome browser plot.

The following Use Cases describe two ELMER runs for the same TCGA Breast Cancer dataset, one using Unsupervised mode, and the other using Supervised. The full HTML reports for these runs, including all source code used, output tables, and plots, can be downloaded here: https://github.com/tiagochst/ELMER_supplemental/raw/master/supplemental_files.zip.

### Use Case 1: Breast Invasive Carcinoma (Unsupervised mode)

We performed *ELMER* (v 2.4.3) analysis comparing 778 Breast Invasive Carcinoma (Primary solid tumor) samples to 83 samples of normal tissue adjacent to the tumor. In this use case we wanted to be able to detect non pre-determined molecular subtypes among the tumor samples, so the percentage of samples used to identify the differentially methylated probes in function *get.diff.meth* was set to 20% and the mode in function *get.pair* and in function *get.TFs* which was set to “unsupervised”. In this mode we define the *U* (unmethylated) group as the samples with lowest quintile of DNA methylation levels and the *M* (methylated) group as the highest quintile.

This analysis showed that the set of hypomethylated CpG probes (DMCs) in the tumors and linked to the expression of a nearby gene (Supplementary Fig. 12) had an enrichment for TFBS motifs for FOX family transcription factors (*FOXA2, FOXA3, FOXA1*, etc.) (Supplementary Fig. 13). For the most highly enriched motif *FOXA2*, the master regulator analysis identified *FOXA1* as the top candidate among all TFs in the human genome (Supplementary Fig. 14), with the collaborating factors *GATA3* and *ESR1* as the next best candidates (Supplementary Fig. 15). This illustrates the important point that *in vitro* defined motifs from public TFBS databases are not always bound by the same TF family member *in vivo*. This was the same as the results from Yao et al. (2015), where we showed that ELMER identification of *FOXA1, GATA3*, and *ESR1* were driven specifically by the ER+ (luminal A and luminal B) tumors. However, our unsupervised analysis (both in Yao et al. and here) did not reveal Master Regulators for the other Breast Cancer molecular subtypes, such as Basal-like, HER2+, etc.

**Supplementary Fig. 12.**
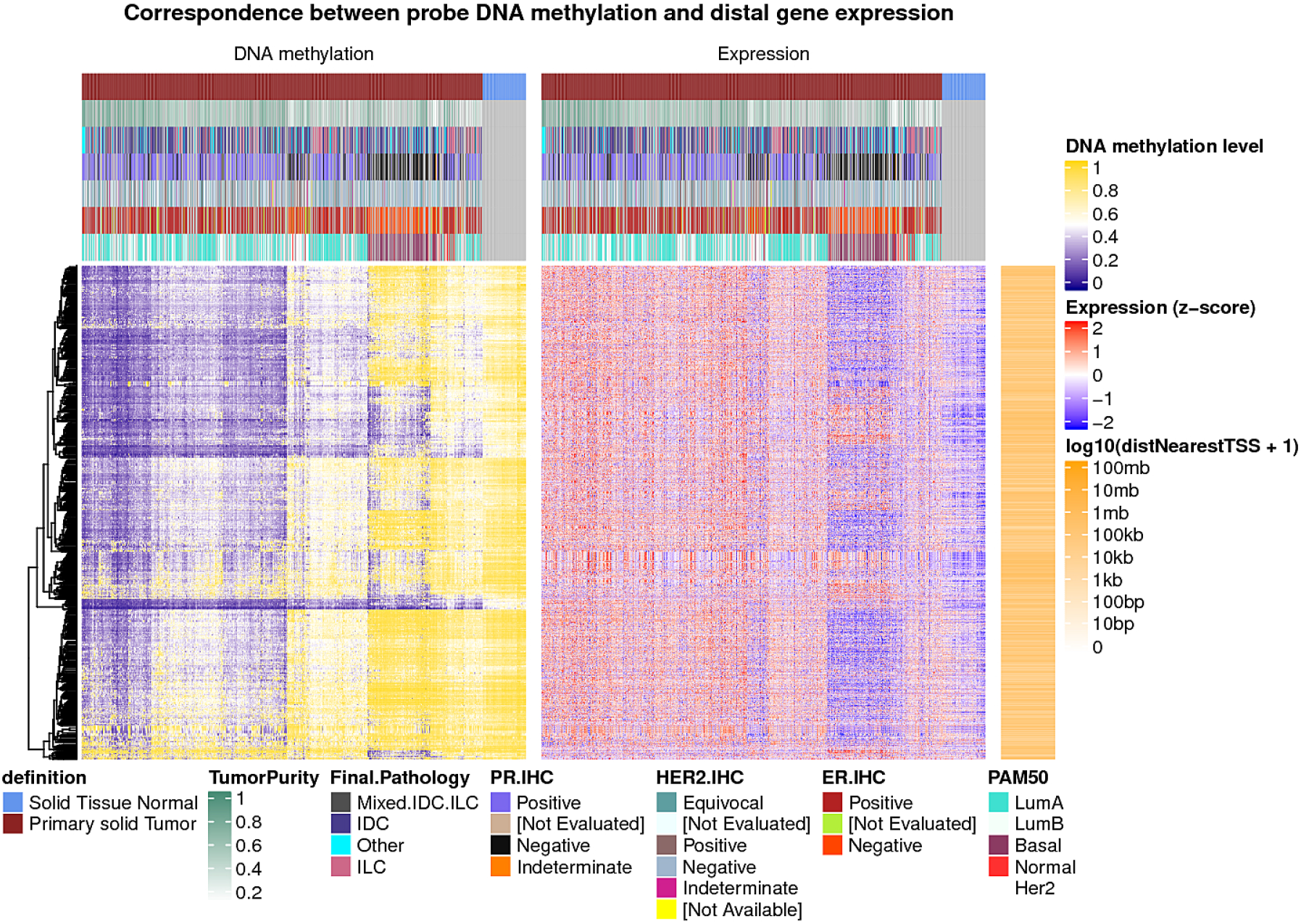
The comprehensive heatmap view shows all probe / gene pairs identified by ELMER, clustered according to similarity. This plot is based on the Unsupervised analysis of Breast Invasive Carcinoma (Primary solid tumor) samples to 83 samples of normal tissue adjacent to the tumor (Solid Tissue Normal). The inverse correlation between methylation and expression can be observed.

**Supplementary Fig. 13.**
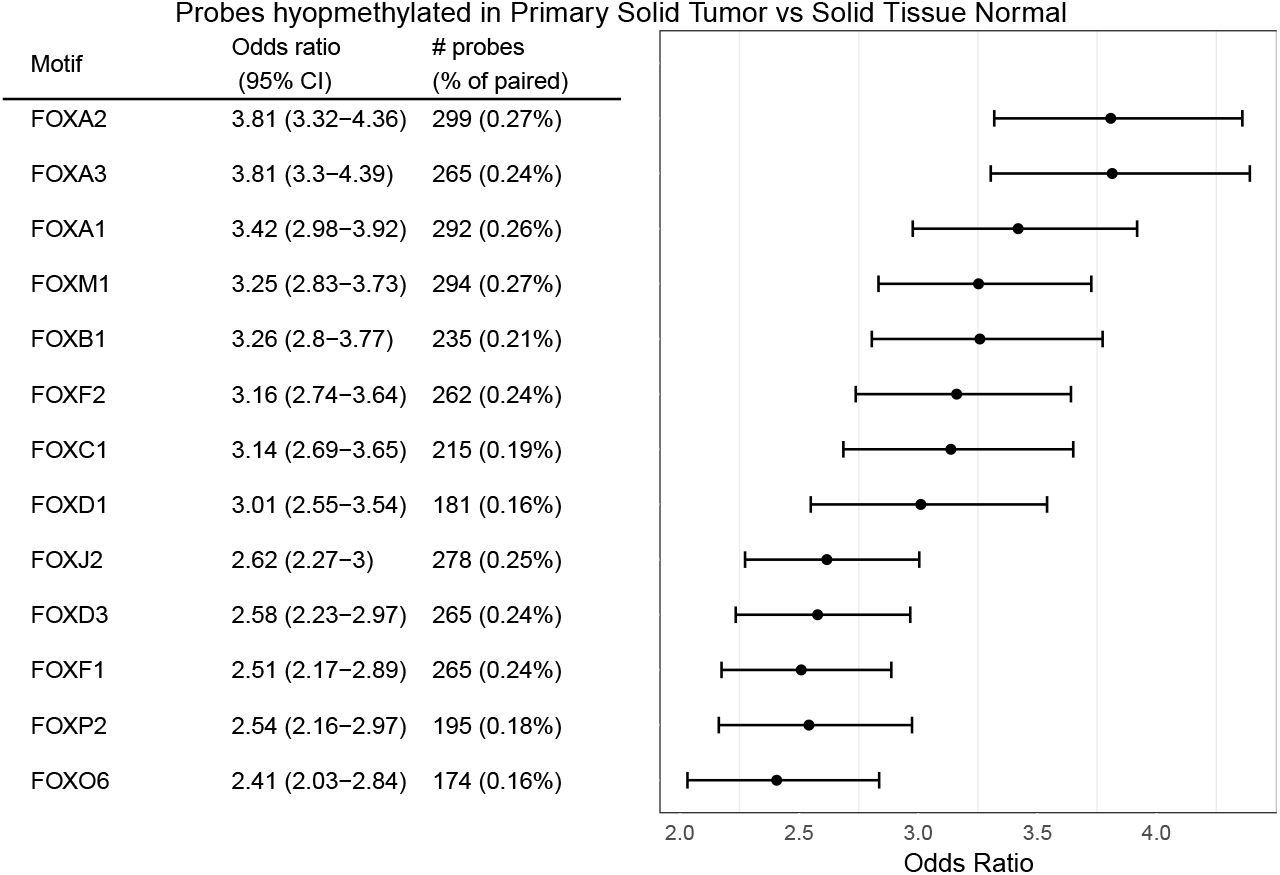
Motif enrichment plot shows the enrichment levels (*OR* ≥ 2.0) for the most significant motifs based on the TCGA Breast Cancer Unsupervised analysis. A number of less significant motifs meet our default OR threshold of 1.1 (*lower.or* = 1.1), which can be browsed in our full Supplemental output report.

**Supplementary Fig. 14.**
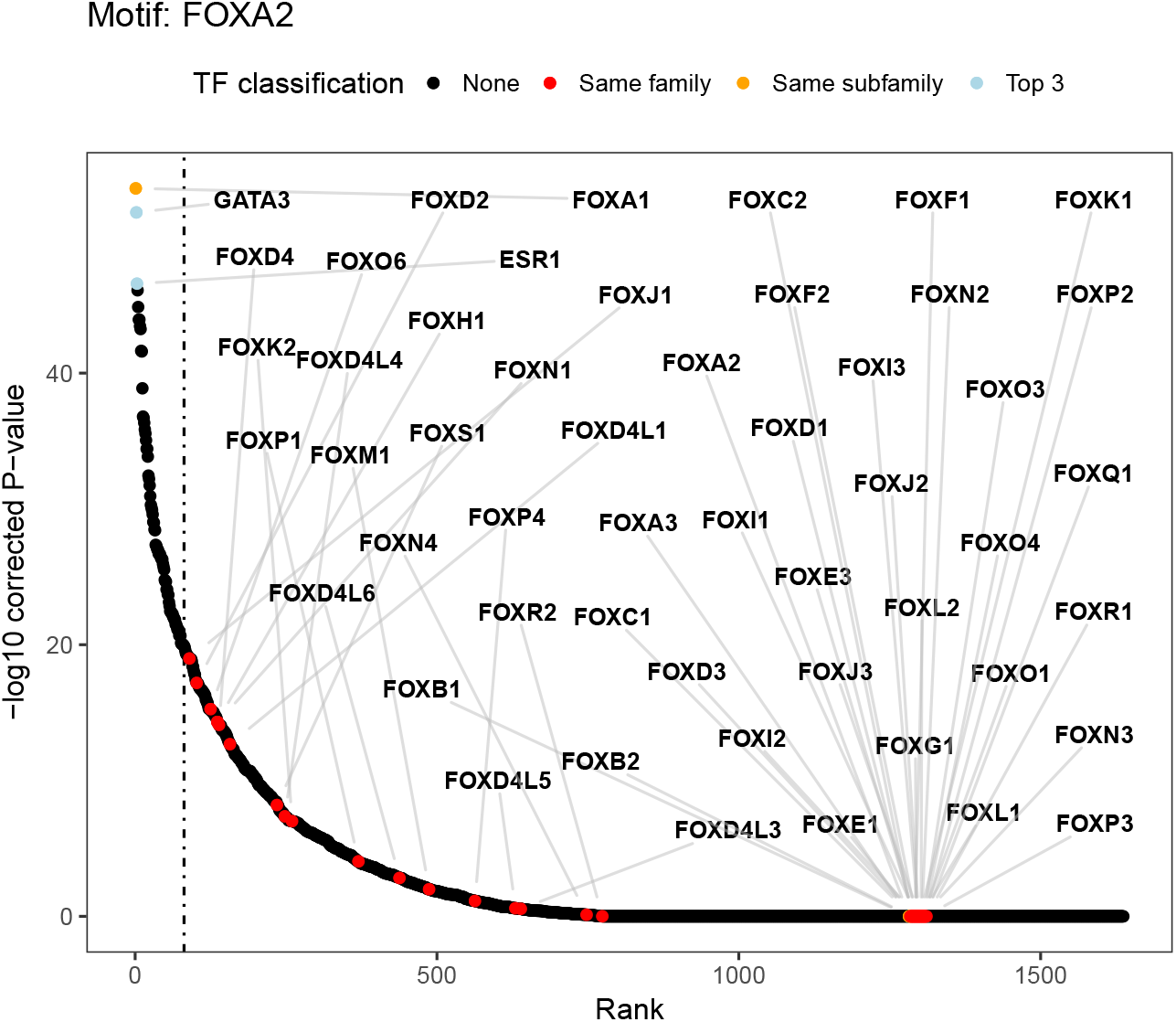
TF ranking plot: For a given enriched motif, all human TF are ranked by the statistical −log_10_(*P − value*) assessing the anti-correlation level of candidate Master Regulator TF expression with average DNA methylation level for sites with the given motif. As a result, the most anti-correlated TFs will be ranked in the first positions. For example, the figure shows the TFs ranking for the (*FOXA2*) motif in which the TF *FOXA1* is ranked in the first position, meaning it is the most anti-correlated TF according to the statistical test. By default, the top 3 most anticorrelated TFs (*FOXA1, GATA3* and *ESR1*), and all TF classified by TFClass database in the same family (Forkhead box factors) and subfamily (FOXA) are highlighted with colors blue, red and orange, respectively. The complete anti-correlation data for the top three candidates (*FOXA1, GATA3*, and *ESR1*) is expanded in Supplementary Fig. 15

**Supplementary Fig. 15.**
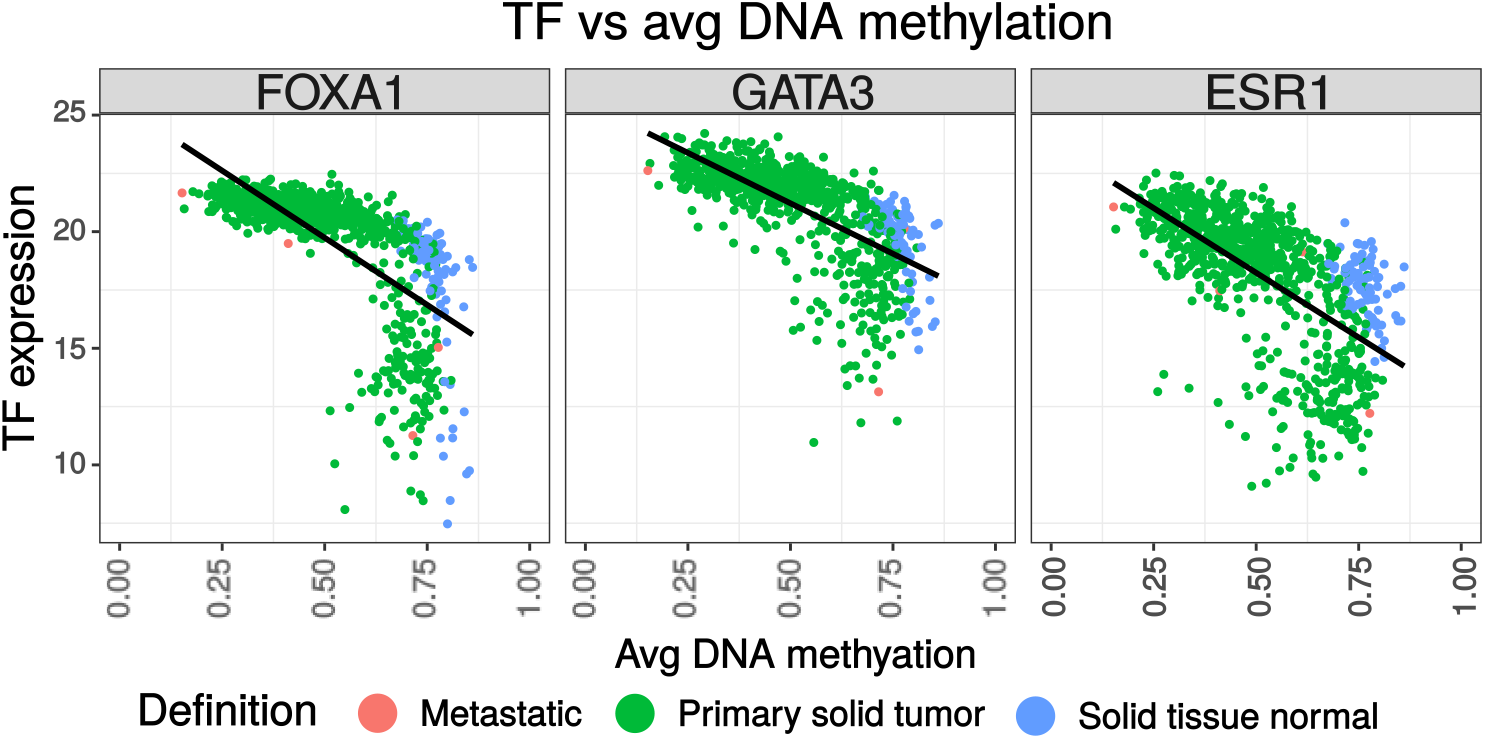
*FOXA1, GATA3* and *ESR1* were identified as the most significant Master Regulator candidates for the top motif (*FOXA2*). All FOX factors belonging to the same TFClass binding family are highlighted.

### Use Case 2: BRCA molecular subtypes analysis (Supervised mode)

Several studies identified distinct molecular Breast Cancer molecular subtypes including luminal-like (Luminal A and Luminal B) subclasses, which are Estrogen receptor-positive (ER-positive), and the basal-like, ErbB2-positive and normal-like subclasses (ER-negative) (Perou et al., 2000; Yersal and Barutca, 2014; S0rlie et al., 2001).

We performed pairwise analysis comparing known molecular subtypes (Her2, Luminal A, Luminal B and Basal-like) using the TCGA BRCA dataset and classifications retrieved from (Ciriello et al., 2015). Supplementary Table S2 shows the number of samples of each molecular subtype of breast cancer and Supplementary Table S3 summarizes the candidate MRs identified.

The *Unsupervised* analysis of the same sample identified several Luminal type Master Regulators (MRs) such as *FOXA1, GATA3*, and *ESR1*. In order to identify MRs for the other subtypes, we created a table (Supplementary Table S3) of candidate MRs identified by each pairwise ELMER run (complete results can be found in the supplemental HTML file described in the Supplementary Methods section).

Interestingly, several new MRs are identified for the Basal-like group, and these were mostly consistent in comparisons against Luminal and HER2+ subtypes. One group of MRs identified are the *SOX10* and *SOX9* TF signatures. For these signatures, the regulatory TF candidate identified are the *SOX9* (Sry-related HMG box-9) TF and *SOX11* (Sry-related HMG box-11) TF; this correlation between basal-like and *SOX11* was recently described by Shepherd et al. and *SOX9* was described by (Gong et al., 2015). Most interestingly, we found *KLF5* to be a consistently predicted MR for the Basal-like breast subtype. *KLF5* is a master pluripotency factor of embryonic stem cells, and has been associated with a number of different cancers. In breast cancer, its overexpression has been linked to aggressive, ER-negative and basal-like breast cancers (Ben-Porath et al., 2008).

**Table S2.** Number of samples of the molecular subtypes of breast cancer

| Molecular subtype | Number of samples |
| --- | --- |
| Basal | 85 |
| Her2 | 34 |
| LumA | 288 |
| LumB | 117 |
| Normal-like | 22 |

**Table S3.**
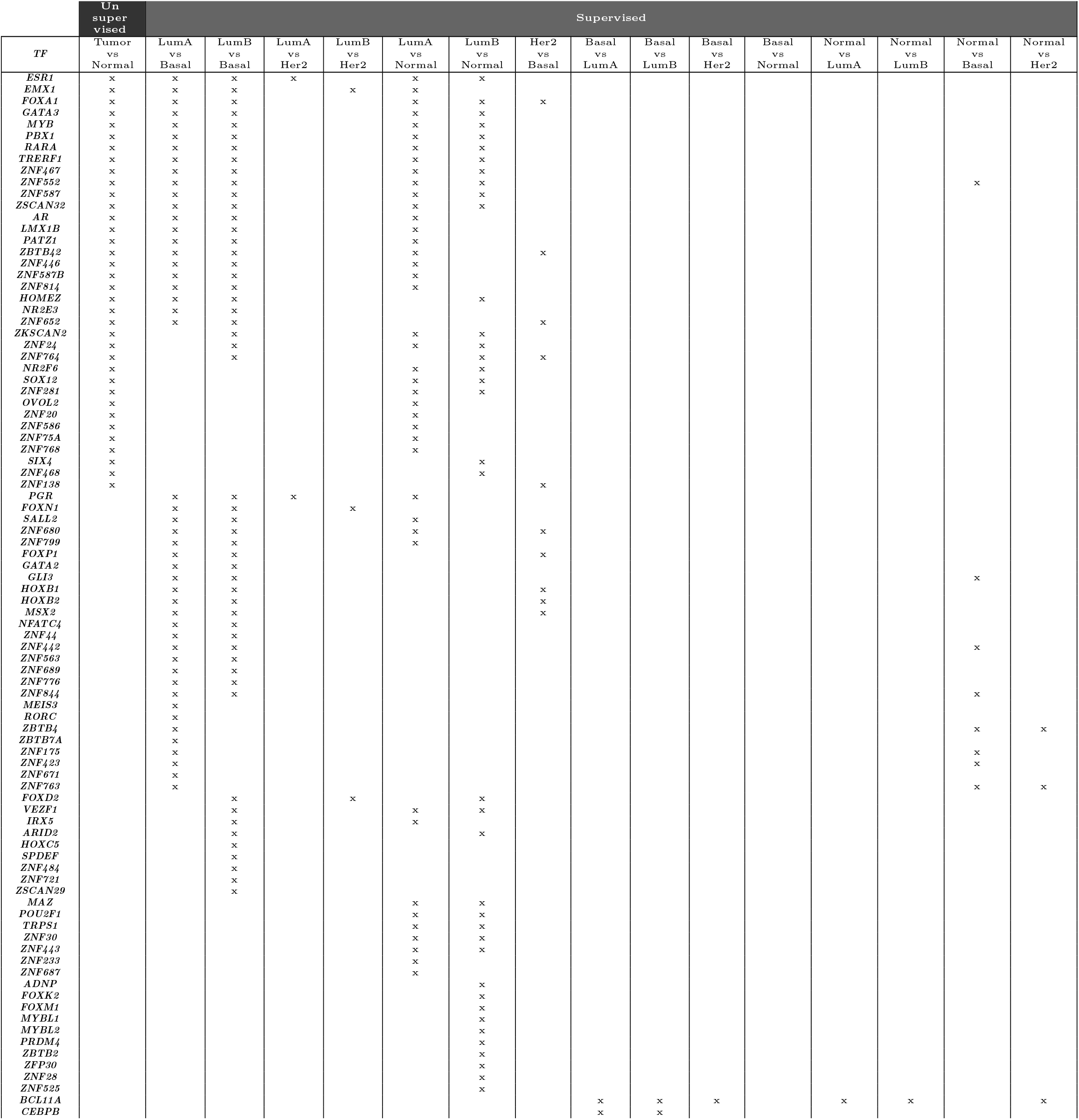
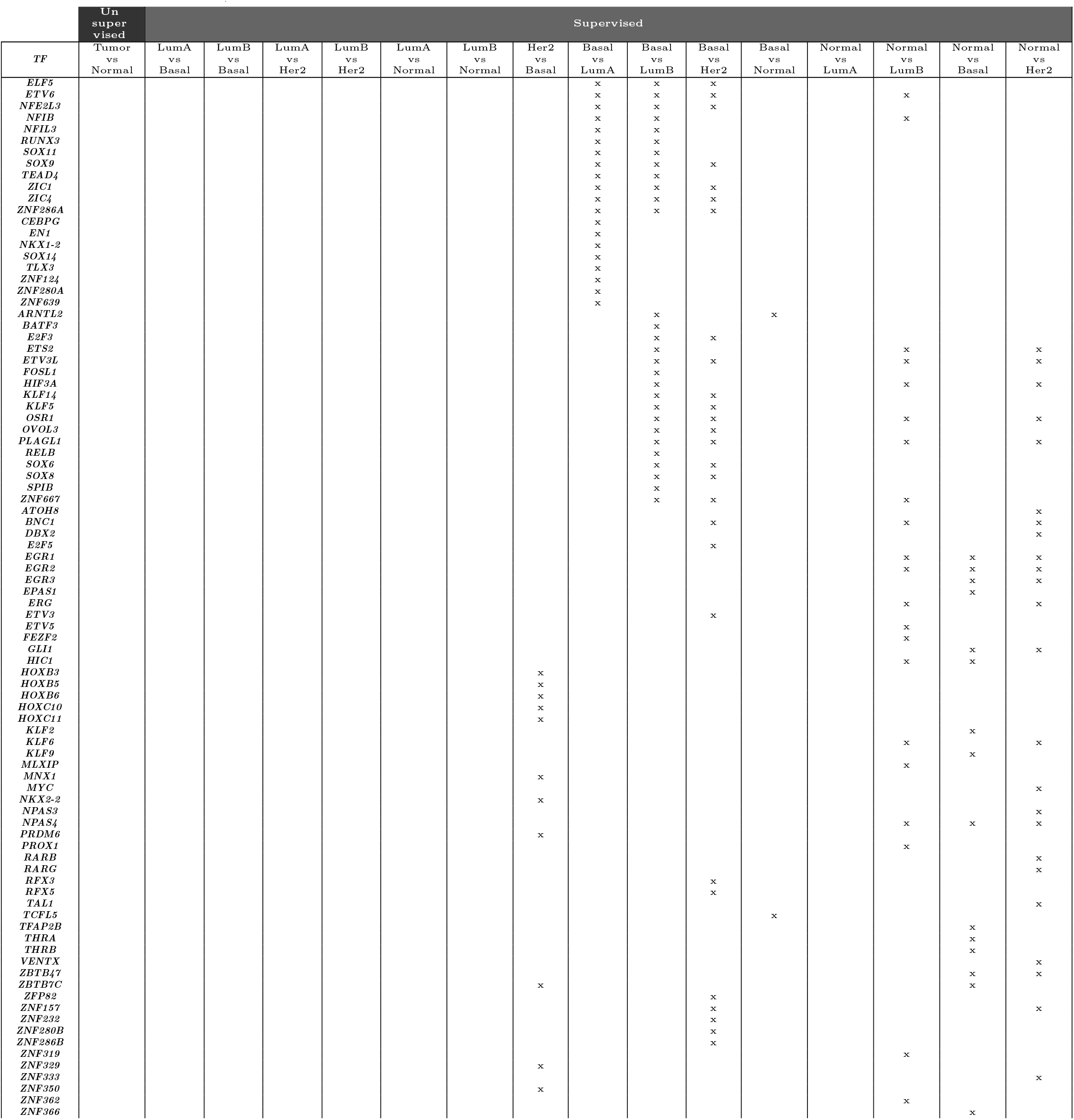
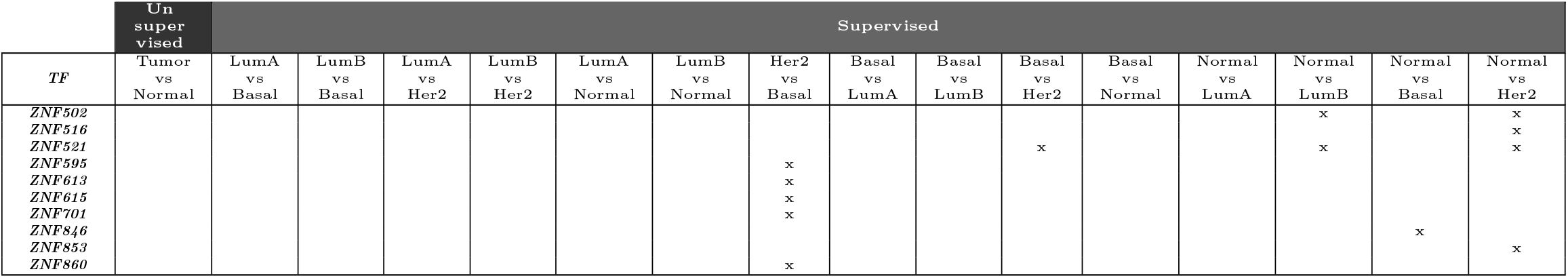
Candidate master regulator TFs (MRs) identified in the supervised analysis (Tumor vs Normal) and unsupervised analysis (pairwise comparison between molecular subtypes: LumA, LumB,Her2, Normal-like,Basal-like). Each column shows a pairwise analysis, identifying the MRs active in the first group. TFs were ordered by the first analysis column where they appear, then by the second one, etc.

### Software engineering best practices

To improve the software error handling we included in this new version: i) unit testing which ensures that our tool is working as expected and any modification in the code does introduce bugs, ii) TryCatch blocks to handle exceptions which will provide user with more information in case a exception is reached, iii) Continuous Integration (CI) services such as Travis (https://travis-ci.org/tiagochst/ELMER) and Appveyor (https://ci.appveyor.com/project/tiagochst/elmer), which not only ensures our tool is installable, free of bugs, passes unit tests and that its documentation can be created after any code modification, but also reduces the time to identify possible platform specific problems. iv) the documentation has been revised and improved by changing the format of a PDF presentation to an HTML-navigable page (Supplementary Fig. 16), v) The data package now has all the code required to easily create all auxiliary objects from publicly available databases, which was not available for ELMER v1.

**Supplementary Fig. 16.**
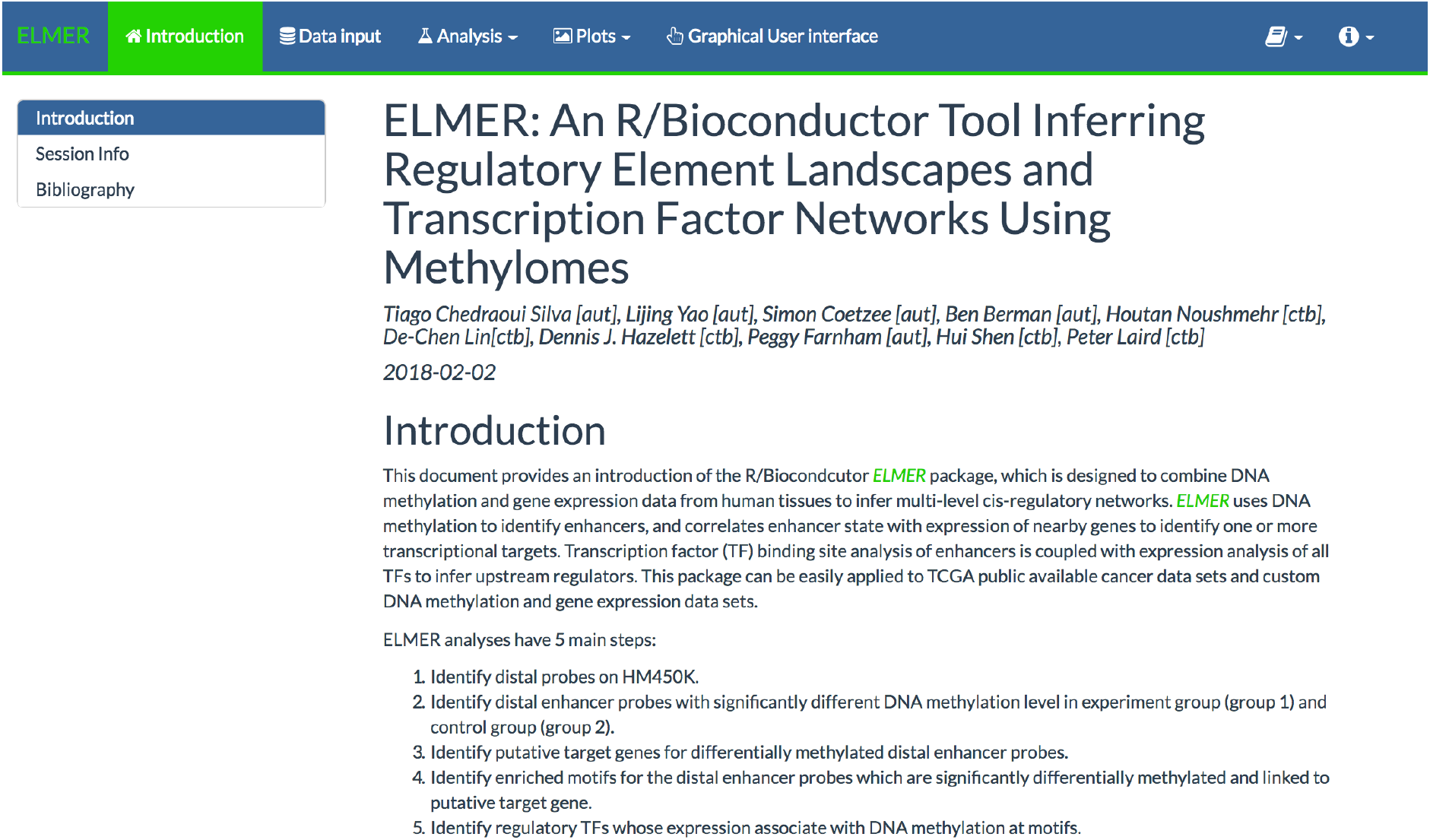
New ELMER documentation available at http://bioconductor.org/packages/devel/bioc/vignettes/ELMER/inst/doc/index.html

### Computational efficiency

To compare the new version with the old one (ELMER v1 vs ELMER v2), we performed the *Unsupervised* analysis of the TCGA breast cancer data set in an Ubuntu 16.04.3 LTS, 32Gb Intel Precision Tower 5810 Intel (R) Xeon (R) of RAM, CPU E5-1650 v3 @ 3.50GHz, using 10 cores for code parallelization. The times for each one of the main functions are shown in Supplementary Table S4.

Some functions had an increase in the time due to changes either in the data or method. As the number of TF binding models used in this new version increased from 91 to 771 it was expected that the function *get.TFs* would increase the time to run, as more iterations will be performed. Also, the enriched.motif now performs and Fisher’s exact test for each motif increasing the time to execute the function. Overall ELMER v2 decrease 55% the time to run the analysis compared to ELMER v1. The code used to run ELMER v2 is the same provided in the HTML reports and the code to run ELMER v1 can be found in this gist (https://gist.github.com/tiagochst/04c2c61b1f3f34f892cd0d0e12a81be6).

Also, Supplementary Table S5 shows the time required to run each ELMER *Supervised* analysis. Only runs that executed all functions were included, that means, if a analysis was not able to identify differential methylated probes it was excluded from the table. Although the larger the number of samples resulted in longer execution time, it is worth remembering that the unsupervised mode uses all samples in all the steps, while the supervised mode uses only a quintile of samples in each group which will reduce its run time.

**Table S4.** Performance comparison between ELMER v1 vs ELMER v2. All values shown are in seconds.

| Function | Time elapsed ELMERv1 | Time elapsed ELMERv2 |
| --- | --- | --- |
| *get.diff.meth* | 163 s | 52 s |
| *GetNearGenes* | 174 s | 267 s |
| *get.pair* | 29468 s | 12790 s |
| *get.enriched.motif* | 2 s | 23 s |
| *get.TFs* | 110 s | 301 s |
| All functions | 29917 s (≈ 8h18min) | 13433 s (≈ 3h45min) |

**Table S5.**
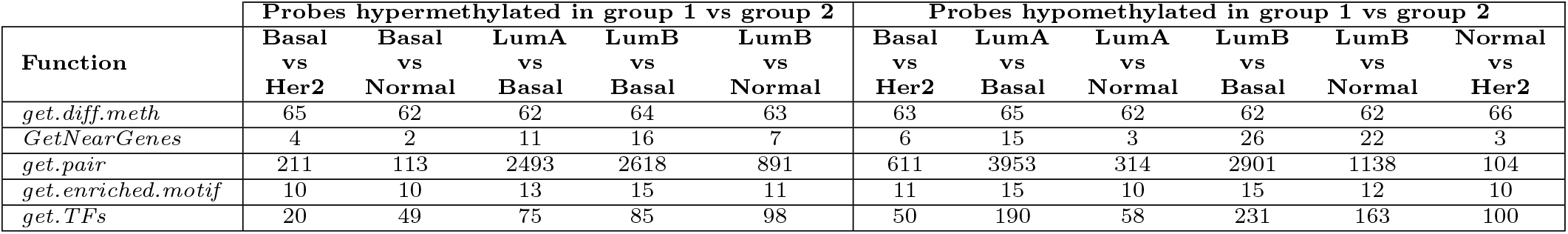
Performance comparison between each supervised analysis using ELMER v2. All values shown are in seconds.

